# Spatial transcriptomics and genetically implicated genes identify putative causal tissue structures for complex traits

**DOI:** 10.1101/2025.05.02.651876

**Authors:** Linda Kvastad, Aaron Kollotzek, Rasool Saghaleyni, Chih-Fan Chang, Tianze Cao, Tuuli Lappalainen

## Abstract

Spatially resolved transcriptomics is transforming our understanding of cellular and molecular diversity of tissues. Here, to identify tissue structures that are enriched for putatively causal disease processes, we integrated 31 human and mouse spatial datasets from 8 organs with genes that are genetically implicated in 32 human diseases. Applying our novel approach STEAM, we identified both known and novel roles of tissue structures in human diseases and identified spatially clustered disease gene subsets. For example, in the brain we observed enrichment of neuropsychiatric disease in cortical layers, and immune and barrier dysfunction in Alzheimer’s disease, with integration with single-cell data highlighting the complementary insights from the two data types. Spatial coexpression of drug target genes with genetically implicated genes in enriched tissue structures showed potential for spatially informed drug repurposing. Altogether, we show vast potential for integration of genetic discoveries with growing spatial datasets to understand human disease biology.

---

Spatially resolved transcriptomics (SRT) is a rapidly expanding field that has provided unprecedented insights into cellular and molecular variation within tissues, advancing our understanding of tissue structures and their biological functions^1,2^.

While much of the field has focused on identifying structures, annotating, and integrating SRT datasets themselves, future opportunities lie in interrogating the biological roles of the identified tissue structures. One key approach to this is gene set enrichment analysis, which asks whether a set of genes – such as genes implicated in a particular disease – are enriched in SRT datasets.

Genome-wide association studies (GWAS) have been critical for identifying genetic variants associated with complex traits and diseases^3^. As genetic data is effective in pinpointing putatively *causal* disease processes - rather than pathophysiological processes caused by the disease or its symptoms - they are being increasingly leveraged in drug development. Indeed, it has been shown that selecting genetically supported drug targets increases new drugs’ clinical development success rate^4–6^. This has inspired major efforts, including the Open Targets platform^7,8^ that provides a systematic resource listing genes implicated in diverse traits. However, further functional studies are needed to understand the underlying causal biological mechanisms. GWAS data has been combined with bulk and single-cell RNA sequencing (RNA-seq) and epigenomic marks to identify enrichments of GWAS signal in different tissues and cell types^9–21^. SRT datasets provide emerging opportunities to assess GWAS enrichment across tissue structures in multiple organs, traits and datasets^22^. Previous studies have mainly focused on the direct use of GWAS summary statistics. Here, we utilize the Open Targets platform^7,8^, where data from multiple GWAS are distilled into trait-specific ranked gene lists. We then use spatially resolved gene expression from healthy tissues to pinpoint the annotated structures where disease-implicated genes are enriched, implicating these structures as potential sites of casual disease processes.

However, such analysis is complicated by the specific characteristics of SRT datasets. Most notably, it is known to be sparse, and the number of genes and unique molecular identifiers (UMIs) are often unevenly distributed. To overcome these challenges, we developed an approach called **S**patial **T**rait **E**nrichment **A**nalysis with per**M**utation testing (STEAM), a robust and scalable computational approach to measure the enrichment of average gene expression across clusters in a spatial dataset from a given gene list; it calculates a permutation p-value and performs multiple testing corrections based on the number of clusters. For ranked gene lists, STEAM can set an empirically derived threshold beyond which the enrichment signal falls to baseline. Thus, STEAM can perform robust gene set enrichment analysis of spatial structures in SRT datasets.

In this study, we analyzed spatial enrichment of genetically implicated gene sets for 32 complex traits and diseases from Open Targets, and combined these data with 31 previously published SRT datasets from eight organs. Applying STEAM to pinpoint putative causal tissue structures for complex diseases, we discovered numerous tissue structure enrichments. For example, our results for Alzheimer disease (ALZ) imply mechanisms beyond amyloid and tau pathology in barrier and immune dysfunction. Furthermore, we analyzed spatial enrichment of drug target genes for these diseases, providing insights into their overlap with genetic evidence and potentially advancing future drug repurposing efforts.

## Results

### Spatial trait enrichment analysis with permutation testing

We analyzed 31 SRT datasets, spanning eight organs^23–35^ (Supplementary Table 1). The spatial structures of each dataset were determined using standard clustering approaches, which included non-negative matrix factorization to identify spatial gene expression patterns (Supplementary Fig. 1-4, Supplementary Table 2-3). We analyzed two biological replicates from humans and/or mice in most organs, and included multiple developmental time points for the heart, intestine, pancreas, and spinal cord. For each tissue we selected the SRT dataset of best quality as an index dataset, treating others as replicates with results shown in supplementary materials, except for the developmental stages where only one sample per stage was included.

We used the Open Targets platform to compile lists of genetically associated and drug target genes for 32 complex traits. Many genes are targets for drugs used in multiple diseases, and some genes are implicated in multiple diseases, but we observed low sharing between genetically associated and drug target genes (Supplementary Fig. 5). With our main focus being enrichment in structures within tissues, we then used previously implicated links between traits and tissues to select trait-tissue pairs for enrichment analysis^8^ (Figure 1a, Supplementary Table 4).

**Figure 1.**
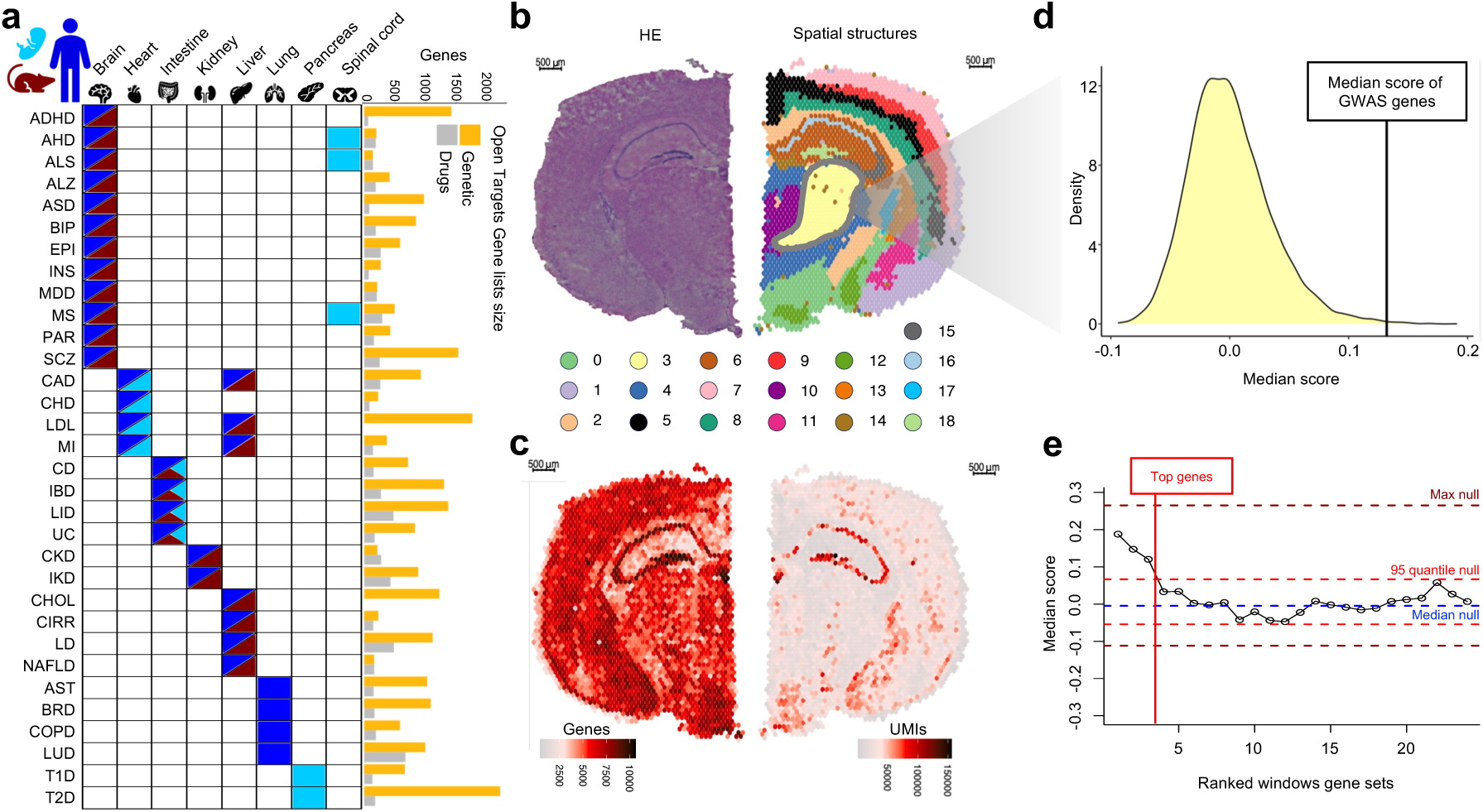
Illustration of the approach and the STEAM pipeline. **a)** Selected trait-tissue pairs, spatial datasets and gene sets included in the study. **b)** Hematoxylin and Eosin (HE) stained mouse brain coronal tissue section (left) and clusters from SRT data (right). **c)** Spatial distribution of number of genes (left) and unique molecular identifiers (UMIs) (right). **d)** Null distribution of the median score for genetically implicated Alzheimer’s genes for cluster 3 in (b). **e)** Median scores for ALZ cluster 3 in (b) in a sliding window of 50 genes, compared to null distribution ranges marked with dotted lines. ADHD=attention deficit hyperactivity disorder. AHD= anterior horn disorder. ALS=amyotrophic lateral sclerosis. ALZ=Alzheimer disease. ASD=autism spectrum disorder. AST=asthma. BIP=bipolar disorder. BRD=bronchial disease. CAD=coronary artery disease. CD=Crohn’s diseases. CHD=congenital heart disease. CHOL=total cholesterol measurement. CIRR=cirrhosis of liver. CKD=chronic kidney disease. COPD=chronic obstructive pulmonary disease. EPI=epilepsy. IBD=inflammatory bowel disease. IKD=inherited kidney disorder. INS=insomnia. LD= liver disease. LDL=low density lipoprotein cholesterol measurement. LID=large intestine disorder. LUD=lung disease. MDD=major depressive disorder. MI=myocardial infarction. MS=multiple sclerosis. NAFLD=non-alcoholic fatty liver disease. PAR=parkinson’s disease. SCZ=schizophrenia. T1D= type 1 diabetes mellitus. T2D= type 2 diabetes mellitus. UC=ulcerative colitis.

To test whether the median expression level of each gene set is higher than the null in the spatial structures of each SRT dataset, we applied our new approach, STEAM. For a robust comparison, STEAM considers the uneven data distribution of genes and transcripts (Supplementary Fig. 6) by creating null gene sets of the same sizes as the trait gene lists studied and performing 10,000 permutations to calculate a p-value for each tissue structure. We applied multiple testing corrections within each disease-SRT dataset pair, and all the reported p-values are after Bonferroni correction based on the number of spatial structures in a given SRT dataset (Figure 1b-d). The robustness of the enrichment results given the uneven sparsity of SRT datasets was verified by lack of correlation between the number of genes or UMIs and nominal p-values (Supplementary Fig. 7).

In the Open Targets platform^7,8^, many complex traits have up to several hundreds of genetically implicated genes (Figure 1a), with an accompanied ranking score indicating the estimated degree of support for each gene-trait association.

However, it is not clear whether all or an unknown top tier subset of these genes are truly trait-associated. To investigate this empirically, we analyzed spatial trait enrichments with STEAM for the selected trait-tissue pairs using a sliding window approach, scanning each gene list from top to bottom using a window size of 50 genes. For many traits, we observed a clear enrichment for highly ranked genes that then decreased to null. We assigned these genes as “top genes” based on a nominally significant sliding window enrichment for at least one tissue structure above the 0.95-quantile (Figure 1 e).

### Putative causal tissue structures for complex traits

We investigated complex trait enrichments for selected trait-tissue pairs in SRT datasets from eight organs (Figure 1 a) and their structures, consisting of 56,516 spatial positions (Figure 2, Supplementary Fig. 8-17). Supplementary Table 5 provides p-values for all trait-structure pairs, including those cited below in the text. We observed a robust signal of enrichment of all genetically implicated genes in the selected trait-tissue pairs (π₁ = 0.55, Figure 2 b) as well as drug targets (π₁ = 0.69, Figure 2 c). The much lower signal for other trait-tissue pairs (π₁ = 0.1 for all genetically implicated genes, Supplementary Figure 8 c) supported the focus on selected trait-tissue pairings to maintain statistical power.

**Figure 2.**
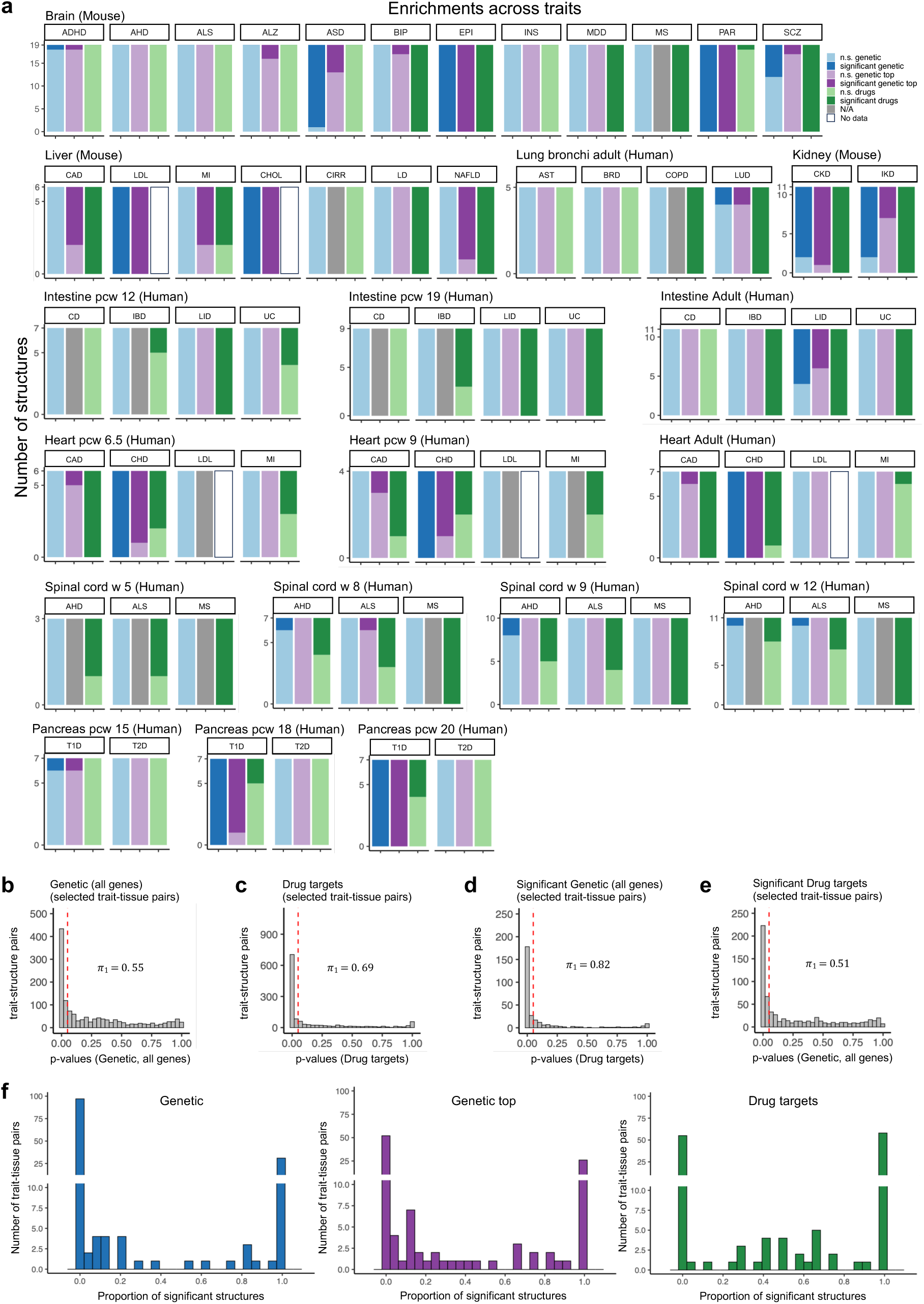
Summary of gene expression enrichments in tissue structures. **a)** Enrichments across traits in all tissues for all genetically implicated genes, top-ranked genetically implicated genes, and drug target genes, visualized for each index sample in each tissue and all included developmental stages. **b-c)** Enrichment nominal p-value distribution of trait-structure pairs, with the proportion of true positives measured as π₁ statistics for **b)** selected trait-tissue pairs for all genetically implicated genes and **c)** selected trait-tissue pairs for drug targets. **d-e)** Sharing of enrichment signal, with **d)** enrichment p-value for drug target genes for trait-structure pairs that were significant for genetically implicated genes, **e)** enrichment p-value for genetically implicated genes for trait-structure pairs that were significant for drug target genes for the same trait. Red dotted line = 0.05. **f)** Proportions of significant structures per selected trait-tissue pair.

Inspecting the proportion of significant structures across tissues for all genetically associated genes, top genes, and drug target genes of selected trait-tissue pairs, we found that in most cases, all or none of the structures were enriched. However, interestingly, we found many trait-tissue pairs (16% to 21% for the three gene classes) where only some structures are significant, pointing to trait enrichments that are specific to tissue structures (Figure 2 f). Examples of these patterns are described below. The signal replicated well between true biological replicates and across species (when data was available, Supplementary Table 5, supplementary figures referred to below).

We next assessed how well current drug targets align with genetic evidence. Structures significantly enriched for genetically implicated genes were often also enriched for drug targets for the same disease (π₁ = 0.82, Figure 2 d), indicating that known drugs target biologically relevant tissue structures. However, the lower proportion of structures enriched for drug targets that are also enriched for genetically implicated gene sets (π₁ = 0.51, Figure 2 e) suggests that drug targets are often expressed also in other tissue structures than those supported by genetics. Similar results were observed when comparing top genes and drug targets (Supplementary Fig. 8 d-g).

### Spatial enrichments of non-brain tissues

For some traits, we observed significant enrichments for genetically implicated genes across the tissue, indicating a broad role of multiple structures in these traits (Supplementary Table 5). Examples of this pattern include adult human heart for congenital heart disease (CHD) (Supplementary Fig. 9 a), liver for low-density lipoprotein cholesterol measurement (LDL), liver for total cholesterol measurement (CHOL), and developmental pancreas during post-conceptual weeks (pcw) 18-20 for type 1 diabetes mellitus (T1D) (Supplementary Fig. 11 e, f).

We also observed cases where a smaller subset of tissue structures showed an enrichment, which highlights the value of spatial data and pinpoints interesting features of disease biology. For example, in the adult heart for coronary artery disease (CAD) we observed enrichment only around the vessels (Supplementary Figure 9 a), which aligns with the known endothelial dysfunction in CAD^36^. For the human dataset of the kidney cortex we observed a broad enrichment for inherited kidney disorder (IKD), in contrast to chronic kidney disease (CKD) where we only saw enrichment in the tissue structures of glomerulus and proximal tubule (p = 0.04; Supplementary Fig. 11 a). In mouse kidney samples both IKD and CKD had cortex enrichments, with CKD having the strongest enrichments in glomerulus and proximal tubule (Supplementary Fig. 11 c). Interestingly, emerging evidence points towards complex interactions between glomeruli and tubules underlying progressive kidney injury, where damage to one tissue structure can cause injury to the other^37^. Our observations for CKD support such a model where two different tissue structures in the same tissue region, here the kidney cortex, could contribute to dysfunction in disease. In the lung, we saw enrichment across the parenchyma tissue for broadly defined lung disease (LUD); this was observed both across the parenchyma datasets and within the bronchi dataset that included a smaller spatial structure of parenchyma tissue (Supplementary Fig. 12 b-d). In the intestine, we observed the strongest enrichment towards the top of the villus for the umbrella category of large intestine disorder (LID) (Supplementary Fig. 13 a), with similar non-significant enrichment patterns in all other datasets (Supplementary Fig. 13 b-e). In addition, using a SRT intestine dataset with a strategy for library preparation that excluded mitochondrial and ribosomal genes we observed enrichments in lymphoid follicles for Crohn’s diseases (CD), and for ulcerative colitis (UC) in lymphoid follicles and part of the submucosa, and part of the submucosa for inflammatory bowel disease (IBD) (Supplementary Fig. 13 b). Similar enrichment patterns in lymphoid follicles were observed for CD in the other adult human and mouse intestine datasets (Supplementary Fig. 13 a and e). CD is known to be an immune mediated disorder with lymphatic dysfunction^38^.

### Spatial enrichments across development

SRT data across developmental time points provides an opportunity to understand how disease-relevant structures manifest during tissue development. For some datasets and diseases, we observed patterns that were generally consistent between time points. For the developmental spinal cord datasets, we observed enrichments for anterior horn disorder (AHD) and amyotrophic lateral sclerosis (ALS) in the anterior ventral horn gray matter tissue (Supplementary Fig. 14), where others have previously found loss of synapses in the ventral horn for ALS^39^. In the human heart from fetal to adulthood, we analyzed two different human developmental time points and a biopsy from the adult human heart (Supplementary Fig. 15), finding congenital heart disease (CHD) to have the strongest enrichments across the heart muscle tissue and CAD around vessel structures.

We analyzed T1D gene set enrichment across developmental ages in SRT datasets from the human developmental pancreas at 15, 18, and 20 post-conceptual weeks (pcw) (Figure 3, Supplementary Fig. 11 d-f). While T1D is usually diagnosed in older children, its developmental etiology is unclear, and it can arise even early in life^40^. During pcw 15, we only observed significant enrichment for T1D genes in structures containing a mixture of exocrine cells (expressing *CTRB1* and *CTRB2*) and endocrine cells (expressing *INS*) (Figure 3 b, c; Supplementary Table 5). In contrast, during pcw 20, all tissue structures were significantly enriched for T1D, although exocrine and endocrine cell-rich structures had the smallest p-values (Figure 3 b, Supplementary Table 5). We further investigated how different subsets of genes contributed to these trait-specific spatial enrichment patterns. We computed spatial coexpression as Spearman correlation of the expression level per spot between all top-ranked genetically associated genes. While most genes did not cluster in a clear way, we identified a small spatially correlated group of key genes, including *INS*, *CTRB1* and *CTRB2* (Figure 3 d). The correlation and anti-correlation pattern show that during pcw 15 and 18, exocrine cells and endocrine cells were spatially colocated and became spatially separated as development progressed (pcw 20) (Figure 3 d). Interestingly, while T1D has been seen as a mainly endocrine (β-cell) driven autoimmune disease, evidence has been mounting for an important role of the pancreatic exocrine cells^14,41^. Our observations support the importance of both exocrine and endocrine cells in T1D disease and exemplify reorganization of disease-relevant cell types during tissue development that can be captured in SRT data.

**Figure 3.**
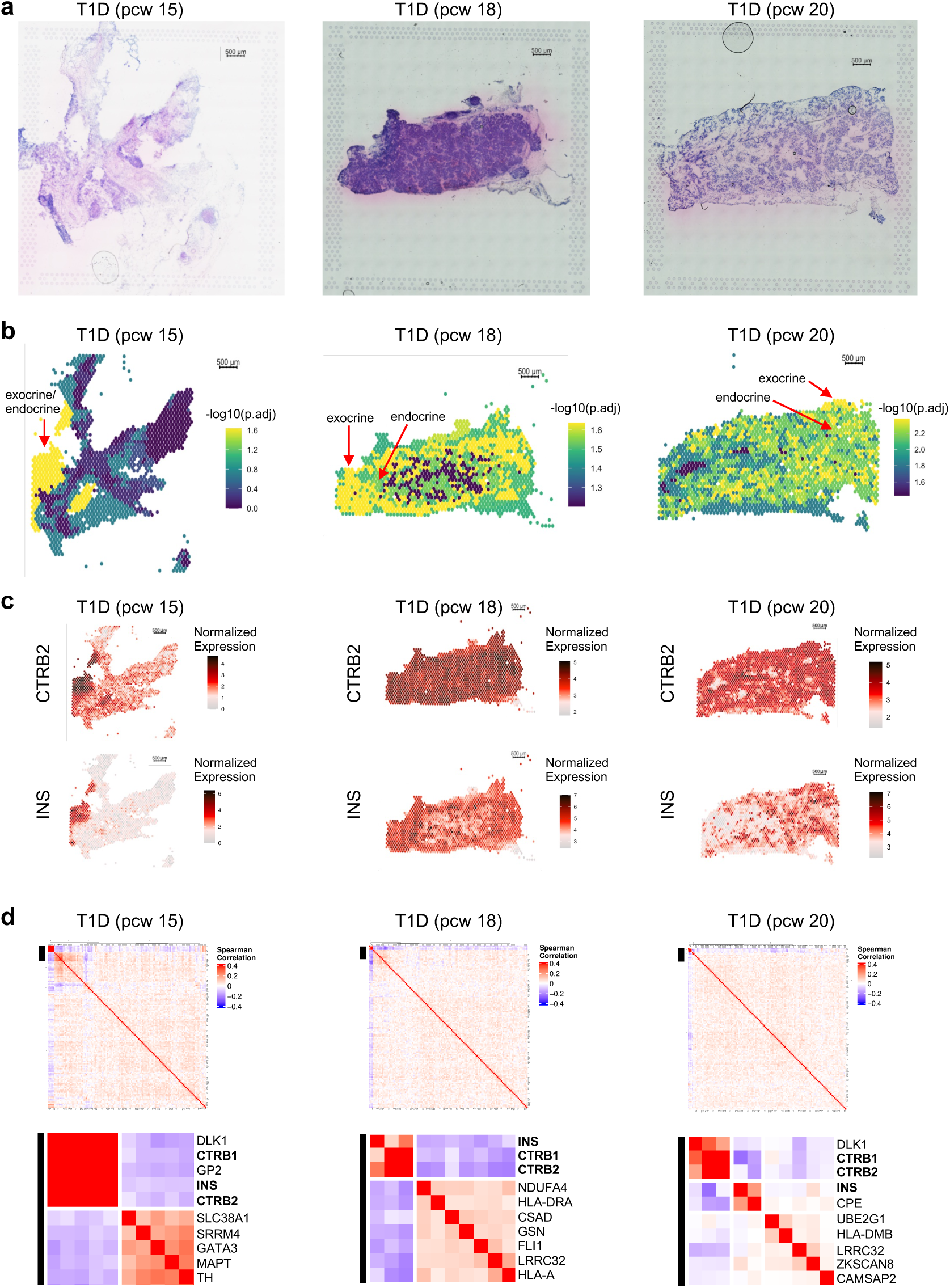
Diabetes type 1 mellitus (T1D) enrichment in the pancreas during human development. **a)** HE-stained pancreas sections for post-conceptual week (pcw) 15 (left), pcw 18 (middle) and pcw 20 (right). **b)** Bonferroni-adjusted p-values for T1D using the top 130 genetically associated genes for the developmental time points. The red arrows show the location of exocrine and endocrine cells. **c)** Expression of selected genes for the time points. **d)** Spatial gene-gene correlation heatmap, including all top genes (top) and zoomed-in top left corner (10 genes) of each heatmap (bottom) for the time points. Post-conceptual weeks (pcw). Scale bar 500µm.

### Spatial and single-cell enrichment of genes implicated in Alzheimer’s and neuropsychiatric disorders

The brain is a highly complex organ with distinct regions performing specific functions. Many neurological and psychiatric disorders have been linked to specific brain regions using neuroimaging approaches^42^, and GWAS enrichment analysis implicating causal cell types^9,12–21^. Here, we investigated spatial enrichment patterns for multiple diseases in the brain (Figure 2 a, Supplementary Fig. 16-17) using the mouse index dataset with the highest quality and coverage of a coronal brain section. In epilepsy (EPI) and Parkinson’s disease (PAR) we observed a broad enrichment across the tissue (Supplementary Fig. 16 a). We found that top-ranked genetically associated genes for ALZ were enriched for choroid plexus, thalamus, and white matter (Figure 4). These findings align with existing literature, which suggests that alterations in the choroid plexus, such as aging-related changes in cerebrospinal fluid dynamics and cytokine levels^43^ play a critical role in ALZ pathology. The thalamus’s involvement in early ALZ stages is reported to be characterized by gray matter decline^44^, and age and severity of dementia with white matter changes^45^, which further supports the relevance of the regions we identified in understanding the disease’s progression. We observed some similarities and interesting differences between mice and human datasets: While comprehensive SRT analysis of the human brain is complicated by its size, both species had significant enrichments in their white matter areas, whereas humans also showed enrichment towards the outer layers of the cortex, in particular layer 1. In addition, in one human sample, we observed the strongest enrichment in spatial structures containing B-cell-related genes, e.g., *IGKC* (Figure 4 a, Supplementary Fig. 18 a, b).

**Figure 4.**
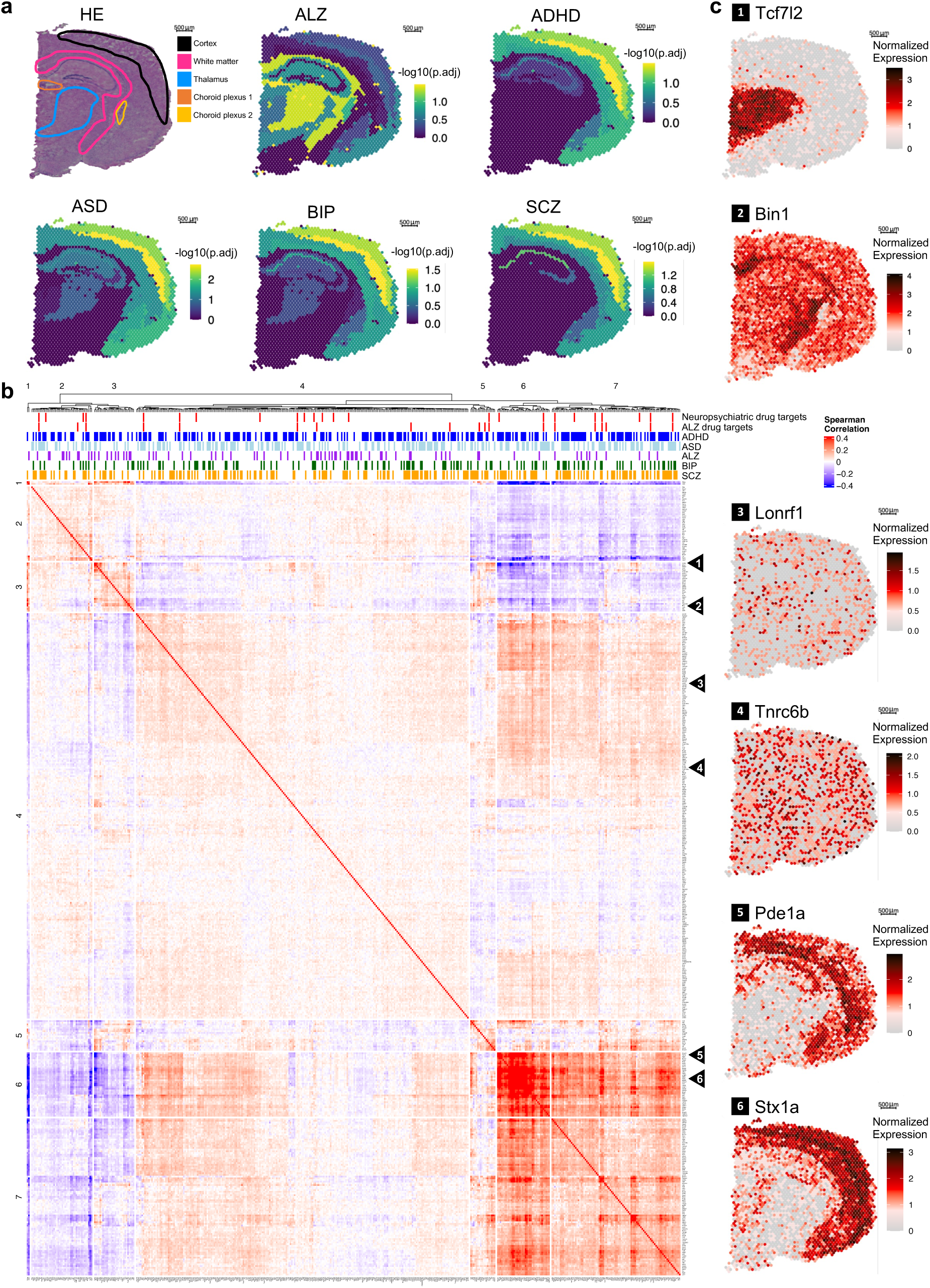
Spatial enrichments of brain related traits. **a)** Mouse brain coronal tissue section with HE staining and annotation of selected regions (top left), and trait-specific Bonferroni adjusted permutation p-values for each spatial cluster, using top genetically associated genes for ALZ (top 70 genes), ADHD (top 200 genes), ASD (top 130 genes), BIP (top 80 genes) and SCZ (top 140 genes). **b)** Spatial gene-gene correlation heatmap, including all top genes for each trait. **c)** Spatial gene expression of selected genes from the heatmap in (b). Scale bar 500µm.

For neuropsychiatric disorders that share parts of their etiology as well as genetic hits^46^ (Supplementary Fig. 18 c), we found that top-ranked genetically associated genes for attention deficit hyperactivity disorder (ADHD), autism spectrum disorder (ASD), bipolar disorder (BIP), and schizophrenia (SCZ) were enriched in the outer layers of the cortex in the mouse dataset (Figure 4 a). In the human dorsolateral prefrontal cortex^24^ we observed a significant enrichment for ASD in layer 2, but not for BIP and SCZ (Supplementary Fig. 17 a). Our results align with previous studies: particularly the ASD enrichment in layer 2/3 has been reported by others^24,47^. While Maynard et al.’s prior pseudo-bulking approach^24^ had a lower resolution than ours, it discovered a more pronounced enrichment pattern for BIP and SCZ. Our observations may be due to data sparsity when analyzing sections individually.

In addition to structure enrichments across the entire gene sets, we further investigated how different subsets of genes contributed to these trait-specific spatial enrichment patterns. We computed spatial coexpression as Spearman correlation of the expression level per spot between all top-ranked genetically associated genes for ALZ, ADHD, ASD, BIP, and SCZ in the mouse index dataset (Figure 4 b-c). Some disease-associated genes clustered into spatially colocalized subsets that may contribute to the disease via distinct mechanisms.

As each structure in a SRT dataset contains a mixture of multiple cell types, we hypothesized that SRT and single-cell RNA-seq (scRNA-seq) datasets provide complementary insights into spatial structures and specific cell types, respectively. To investigate this, we analyzed the Allen Brain Atlas scRNA-seq dataset of the mouse cortex and hippocampus^48^, unfortunately lacking data for the choroid plexus, the thalamus and the white matter that had spatial enrichments in ALZ. Thus, we also analyzed the current most comprehensive human brain single-nuclei RNA-seq (snRNA-seq) dataset^49^. Cell type deconvolution^50^ of the mouse SRT dataset showed biologically meaningful structure-specific cell type proportions (Figure 5 a, b). Next, we applied STEAM to the sc/snRNA-seq datasets to investigate the cell type enrichments of genetically associated genes. To decrease the sparsity of datasets we applied a meta cell approach (see methods), and to empirically redefine the genetic top genes we used the sliding window approach (Supplementary Fig. 19 - 23). Supplementary Table 6 and 7 contain all the p-values for cell type - trait pairs described below in the text. We also did a complementary enrichment analysis of genetically associated genes among the genes that were upregulated and differentially expressed genes (DEGs) in each cell type, using Fisher’s exact test (Figure 5 c, d; Supplementary Fig. 24 and 25).

**Figure 5.**
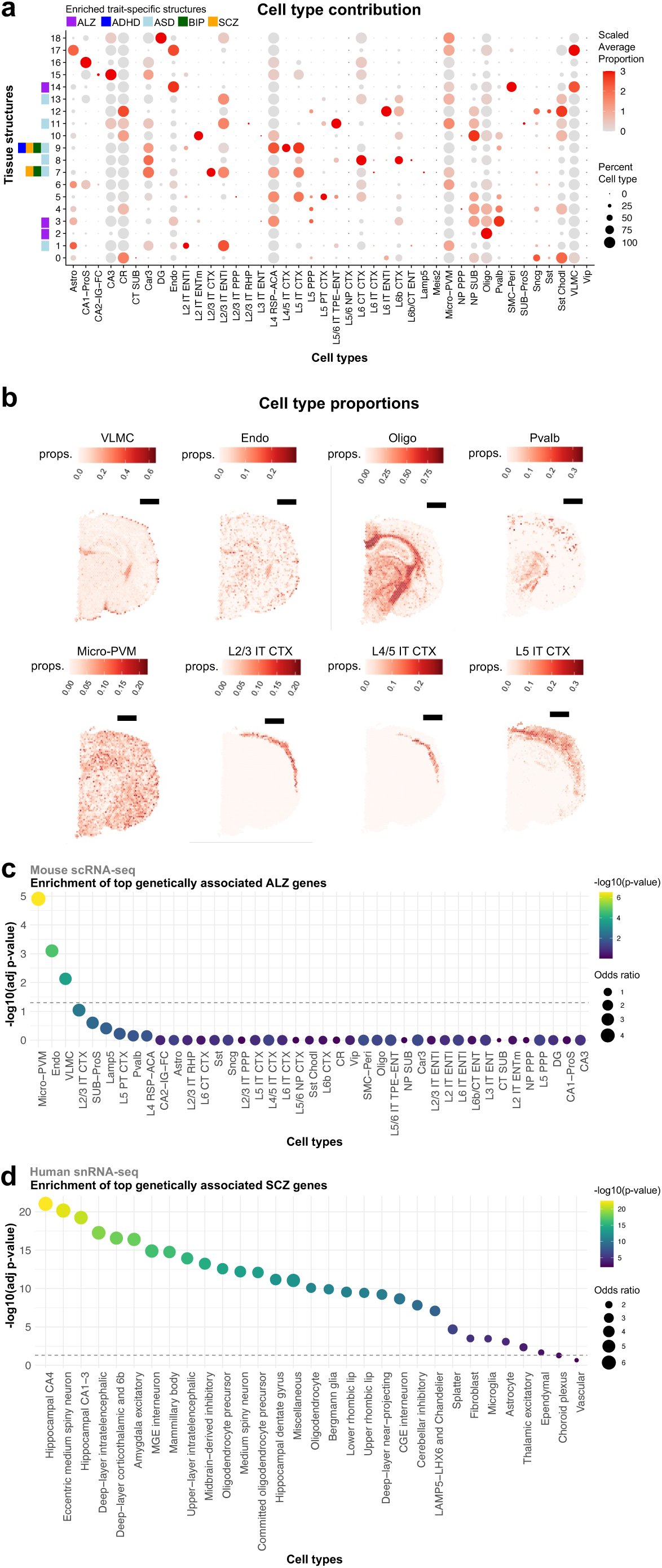
Single-cell contribution to spatial enrichments in the brain. **a)** Cell type composition of the tissue structures of the index mouse coronal tissue section (Figure 1 b, Supplementary Figure 1 a), using the Allen Brain Atlas of the mouse cortex and hippocampus^48^ scRNA-seq data. Enriched spatial structures are marked with a color legend, with **b)** selected cell types shown. Scale bar 1mm. **c-d)** Trait enrichment across cell types for top genetically associated genes among differentially expressed genes **c)** for ALZ across subclasses in the mouse cortex and hippocampus^48^ and **d)** for SCZ across superclusters in the adult human brain^49^. The full list of mouse cell subclass abbreviations can be found in Supplementary Table 6; those shown in **(b)** are: VLMC = Vascular/leptomeningeal cells. Endo = Endothelial cells. Oligo = Oligodendrocytes. Pvalb = Pvalb positive cells. Micro-PVM = Microglia/perivascular macrophages. L2/3 IT CTX = Layer 2/3 intratelencephalic neurons in the isocortex. Human cell supercluster abbreviations; CA = Cornu Ammonis. MGE = Medial Ganglionic Eminence. CGE = Caudal Ganglionic Eminence. Grey dotted line, adj. p-value = 0.05.

For ALZ, in mouse we observed the strongest enrichments of top genes in microglia/perivascular macrophage (Micro-PVM) cells, vascular/leptomeningeal cells (VLMC), endothelial cells, and astrocytes (Supplementary Fig. 19 a), with complementary enrichment in both species showing similar results, with the strongest enrichment in microglia (Figure 5 c, Supplementary Fig. 24 i and 25 a, c). Of these, VLMC and endothelial cells were quite specific to choroid plexus structures that were enriched for ALZ genes (Figure 4 a: choroid plexus 1; Figure 5 a: structure 14). In contrast, Micro-PVM cells showed little structure-specific enhanced contribution and were widely spread across all tissue structures (Figure 5 a and b), indicating potential contributions in disease processes that can take place across the brain. The human snRNA-seq dataset recapitulated many of these patterns, including enrichments in the choroid plexus and vascular cells, and various stages of developing oligodendrocytes (Supplementary Fig. 21 a and 25 a, c). The cell types underlying ALZ enrichment in the white matter and in the thalamus were less clear, but both were enhanced for oligodendrocytes with some ALZ enrichment (structure 2 and 3, respectively in Figure 5 a and b).

For ADHD, ASD, BIP and SCZ, we observed broad significant enrichments of top genes across the mouse neuronal cell types, astrocytes and oligodendrocytes (Supplementary Fig. 19 and 20). In addition, the complementary enrichment showed the strongest signal for the neuronal cell types (Supplementary Fig. 24). The spatial structures enriched for neuropsychiatric disease genes had a high proportion of these cell types, such as L4/5 and L2/3 intratelencephalic neurons in the isocortex (IT CTX) (Figure 5 a, b). In the human snRNA-seq dataset covering cell types from the whole brain, using the complementary analysis of differential expression, e.g., in SCZ the strongest signals from top genes were found in hippocampal cornu ammonis (CA) region CA4, eccentric medium spiny neurons and regions CA1-3 (Figure 5 d), and for all genetically associated genes in committed oligodendrocyte precursor cells (OPCs) and medial ganglionic eminence (MGE) interneurons (Supplementary Fig. 25 i). These findings align with previous literature for SCZ enrichment in medium spiny neurons, hippocampal pyramidal cells, certain interneurons and oligodendrocytes^15,16^, as well as more recent findings of a broad enrichment among neuronal cell types, and MGE interneurons in SCZ^21^. These examples showcase the complementary power of utilizing both SRT and scRNA-seq to investigate cell and tissue enrichments of complex traits. Our observations in mouse and human points towards possible trait contributions from more than one cell type in the same spatial structure.

### Potential for spatially informed drug repurposing strategies

Repurposing approved drugs for new diseases provides an important and increasing line of drug development. We investigated the potential for spatial enrichment patterns to provide additional information to identify repurposing opportunities. Using the brain to demonstrate this principle, we leveraged spatial coexpression of all genes genetically implicated in a given disease (Figure 1 a) to pinpoint those that are expressed in structures identified as disease-relevant (Figure 4) and also targets of known drugs, targeting brain or non-brain traits (Figure 6, Supplementary Fig. 26-28, and Supplementary Table 8). For example, ALZ-implicated gene *Kif5b* is a target for selpercatinib and pralsetinib used in cancer treatment. It was spatially colocalized with genes such as *Bin1* expressed in the white matter, which was overall enriched for ALZ genes (Figure 6 a and 4 c). Interestingly, others have recently reported that a decrease in *KIF5B* levels could slow down and/or prevent abnormal tau behavior in animal and cell models^51^, making it an interesting drug repurposing candidate. In addition, the spatially colocalized ALZ-implicated gene *Clu* is spatially colocalized in the same structures, is also a target for the cancer drug custirsen, and has been suggested to play a role in amyloid beta peptide clearance in an animal model^52^ (Figure 6 a). Voltage-gated potassium and calcium channel genes such as *KCNB1*, *KCNV1*, and *CACNA2D1*, are associated to ADHD, ASD, BIP and SCZ^53–56^ and targets for multiple drugs used in diverse diseases such as multiple sclerosis, gastrointestinal disease, cardiovascular disease, and restless legs syndrome. They were spatially coexpressed with other genes of the cortex that showed a clear overall enrichment for these traits (Figure 6 b, Supplementary Fig. 26-28). Similar spatial gene expression patterns were observed for these genes in the human cortex dataset (Supplementary Fig. 29). This supports the recent rekindled interest in application of these drugs in neuropsychiatric disease^57^. Our observations suggest that considering spatial enrichment of genetically implicated genes could be an interesting additional layer of evidence for identifying potential drug targets and drug delivery locations that intervene with causal processes of disease.

**Figure 6.**
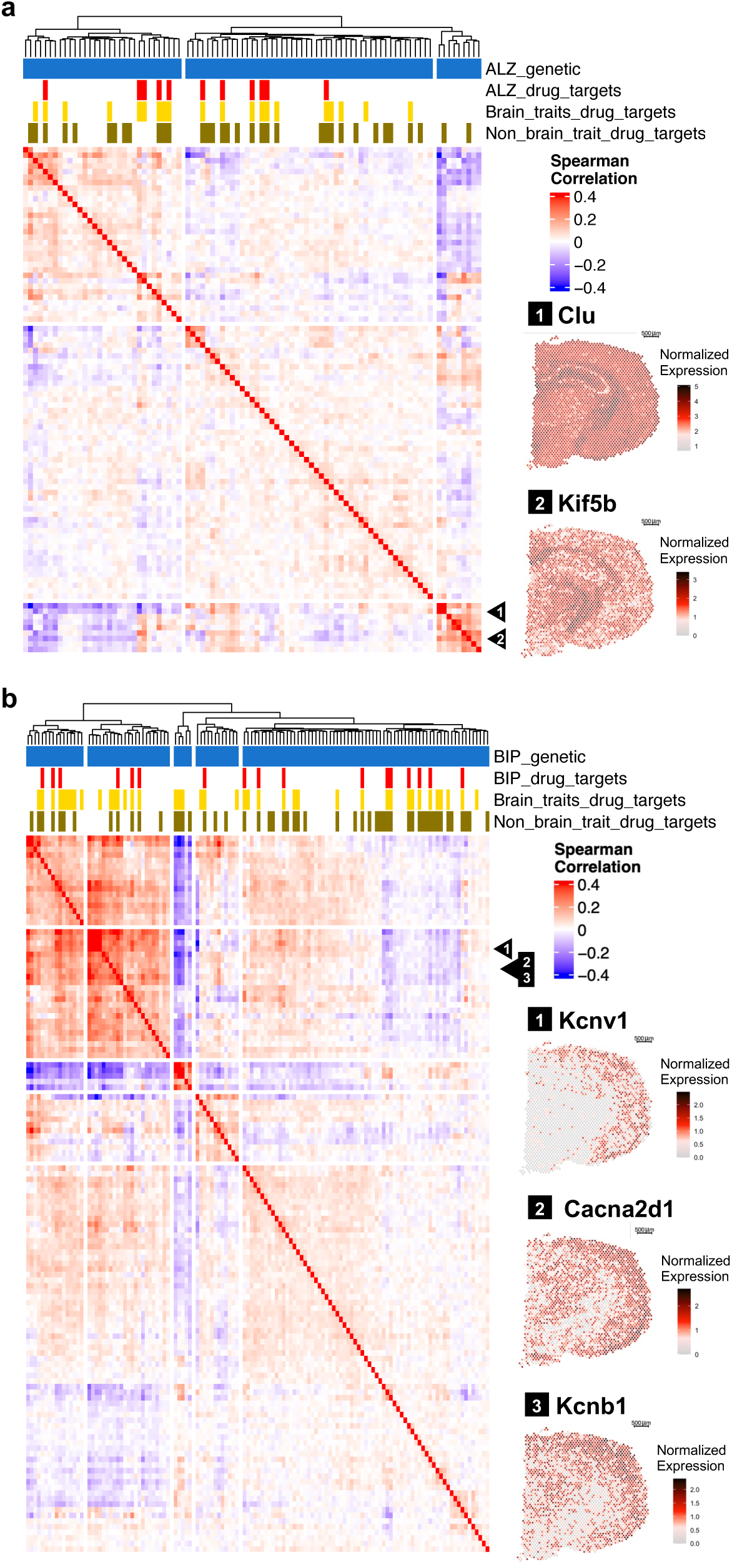
Spatial patterns of drug targets. Gene-gene spatial correlation heatmaps for the index mouse brain coronal tissue section, for genes genetically implicated in **a)** ALZ and **b)** BIP, with their drug target status indicated. Spatial gene expression of selected genes from the heat maps are shown on the side, scale bar 500µm.

## Discussion

Localizing causal disease processes in human tissues is a key part of the quest to understand disease biology and developing interventions. Integrating GWAS data with emerging spatial datasets provides a new opportunity for this. Here, we presented STEAM, a generalizable approach for spatial trait enrichment analysis with permutation testing, and applied it to genetically implicated genes and drug targets for diverse diseases and SRT datasets from a large number of tissues and developmental stages, as well as two supporting brain sc/snRNA-seq datasets. While genes for some diseases are enriched tissue-wide, in up to a fifth of our spatial data we found evidence pointing towards tissue structure-specific trait enrichments, highlighting the value of the spatial resolution. Many of these discoveries were consistent with known biology, but we also identified links between diseases and tissue structures that are not well described in prior literature and indicate potentially interesting targets for further follow-up. While here we focused on 38 disease-tissue pairs based on prior indications, we anticipate that larger data sets with better spatial resolution, replicates, and higher accuracy in linking genes to diseases will further boost the power to allow even broader discovery across all tissues.

Given its refined structures with different biological functions, the brain provides a particularly interesting target for spatial enrichment analysis and further exploration of how spatial enrichments compare to single-cell analysis. Our results show clearly the previously known enrichment of neuropsychiatric disorders ADHD, ASD, BIP, and SCZ in the outer layers of the cortex and cell types such as L4/5 and L2/3 IT CTX that are present in these structures. For ALZ we identified spatial enrichments in part of the ventricles for choroid plexus, thalamus, and white matter in the mouse SRT data. In addition, for the human SRT datasets we observed enrichments in cortical structures, where impaired plasticity has been associated with impaired working memory^58^. While we only observed spatial structures containing B-cell-related genes in the human brain samples, we note that others have shown that B-cell depletion in mice reverses disease progression^59^. The sc/snRNA-seq analysis showed ALZ gene enrichment in VLMC and endothelial cells that are both quite specific to the highly vascularized choroid plexus, surrounding and inside blood vessels and with a role in maintaining barrier integrity^60–62^. On the other hand, the cell type well known for high enrichment of ALZ signal, microglia, were present across multiple brain structures. These results suggest ALZ mechanisms beyond amyloid and tau pathology, involving barrier and immune dysfunction. More generally, these results indicate that some disease processes can arise from dysfunction in a particular cell type that may or may not be spatially enriched, while others may derive from dysfunction of multiple colocalized cell types jointly (and perhaps interactively) in a given tissue structure. Our results for CKD in the kidney even suggests a scenario where multiple distinct structures in a tissue region contribute to dysfunction. Spatial correlation analyses further help to identify subsets of disease genes potentially involved in different disease mechanisms.

In addition to our analysis of genetically implicated genes, we also analyzed spatial enrichments of genes that are drug targets for different diseases. These enrichments often captured the same structures as those discovered via genetics, but drug target genes were enriched more broadly in structures not implicated by genetics. We further used our findings to explore strategies for drug repurposing by identifying situations where a known drug target is not only genetically associated with the disease but also expressed in a structure enriched for genes of that disease, further supporting its biological importance and potential as a repurposing target.

Spatially resolved transcriptomics is still a relatively new field, and many of the limitations of our study arise from the caveats in the current data. Mouse tissues may not fully recapitulate human biology, but the size of human organs limits and complicates their spatial analysis. Differences in enrichment results across intestine datasets highlight the impact of library preparation methods, particularly the exclusion of mitochondrial genes with protein-coding probe-based library preparation. While our approach is planned to be robust to varying gene and UMI count distributions, low-quality datasets will have reduced power. It would be ideal to have single-cell resolution in spatial data rather than analysis and integration of both data types in parallel.

Our approach is based on gene sets compiled by Open Targets – rather than e.g. heritability enrichments of genome annotations^15,16,21^ – as this links more directly to spatial data, facilitates accounting for its properties, and is a natural fit for multi-species analysis. However, linking genetic associations to genes and scoring these is not without problems. We empirically identified the top-ranked gene set that showed a consistent biological signal of spatial enrichment, suggesting that the disease contribution of some of the lower-ranked genes may be questionable. We anticipate that future improvements will allow more accurate gene identification and prioritization and boost the power and resolution of enrichment analyses.

In summary, our study discovered spatial enrichments for a large number of traits across many organs, developmental stages and cell types. Integrating the growing spatial and genetic datasets provided a rich source for biological discovery to characterize mechanisms of human disease.

## Methods

### Spatially resolved transcriptomics datasets

A summary of the datasets’ names and references to their sources can be found in Supplementary Table 1. For each tissue and developmental stage we selected the best quality SRT dataset as an index dataset, as indicated in Supplementary Table 1. Quality was assessed based on the number of genes and UMIs of the dataset, and the HE image field of view including structures of interest. When biological replicates were available, results were shown in supplementary materials. For one SRT dataset (intestine_human_adult_RRST_349_B1) a markedly different 10X library preparation was used, with a transcriptome-wide protein-coding probe-based capture strategy, e.g., excluding mitochondrial and ribosomal genes.

### Spatial transcriptomics data pre-processing

The brain and kidney dataset from 10X Genomics were pre-processed with SpaceRanger (version 1.1.0). Subsequent analyses were carried out using the STUtility^63^ (version 1.1.1) and Seurat^64^ (version 4.3.0) R-packages. Filtering was performed using the *InputFromTable* function in STUtility, with slightly different settings depending on library preparation. For Visium fresh frozen, Visium RRST, and SRT 2k array library preparations, all spots containing less than 500 UMI counts were removed, and genes were removed if they were present in less than 5 spots or had a total UMI count below 100. For the SRT 1k array library preparations, all spots containing less than 500 UMI counts were removed, and genes were removed if they were present in less than 5 spots or had a total UMI count below 20.

### Defining spatial structures

Each dataset was normalized using the *SCTransform* function in Seurat, returning the top 3000 variable genes. Spatial gene expression patterns were identified using the *RunNMF* function in STUtility, the number of factors used and factors included for clustering are stated in Supplementary Table 5 for each dataset. Clustering was performed using the *FindNeighbors* function in Seurat, based on NNMF factors, where factors were excluded if they i) did not show a clear spatial gene expression pattern, ii) were driven mainly by only one gene, iii) were driven by mitochondrial and/or ribosomal genes, or iv) were driven by hemoglobin genes. Then using the *FindClusters* function in Seurat, with individual clustering resolution for each SRT dataset (Supplementary Table 5). To detect differential gene expression, each dataset was first normalized and scaled using the *NormalizeData* and *ScaleData* functions in Seurat. Differentially expressed genes for each seurat cluster were calculated using the *FindAllMarkers* function in Seurat, with custom settings, only.pos = TRUE, and min.pct = 0.25. All the differentially expressed genes (DEGs) for each dataset and cluster can be found in Supplementary Table 3.

### Tissue structure annotation

Annotation of enriched tissue structures was done based on the original publications (Supplementary Table 1). For the 10X mouse kidney dataset we annotated the cortex based on the structural HE similarities of the mouse dataset from Ferreira et al^26^. For the 10X mouse coronal brain dataset we annotated the enriched structures based on: i) Allen Brain Atlas of mouse coronal sections^65^, ii) marker genes from top DEGs of spatial structures associated with specific brain structures according to the Human Protein Atlas^66^ (www.proteinatlas.org), iii) and based on the structural HE similarities and cluster annotations of the coronal mouse dataset from Kleshchevnikov et al^31^. In this article, we interchangeably refer to clusters as structures in spatial datasets and cell types in sc/snRNAseq datasets.

### Open Targets gene lists

For genetically associated genes, the platform uses data source specific association scores and calculates a harmonic sum, which is normalized to create scores between 0 and 1. Evidence is also weighted depending on its source, calibrating its relevance compared to the other data sources^8^. For ChEMBL drug target genes, higher scores are given to targets linked to approved drugs and in a descending order based on their stage in clinical trials, and down-weighted based on failure in clinical trials or severe adverse effects^8^. For the Open Targets versions of each gene list, see Supplementary Table 4.

### Open Targets gene list filtering, pre-processing and sliding windows

Gene lists were filtered for each dataset: if a gene on the list was not present in the dataset (after initial dataset filtering) the gene was removed. All genetically associated gene lists were ranked based on the open target association score, while drug targets gene lists were used as is. For mouse datasets, human gene names were converted using the *useMart* function from the biomaRt^67^ (version 2.52.0) R-package, using Ensemble version 109 from February 2023. For cases where multiple human genes had the same mouse homolog, we only kept the open target gene list data with the highest overall association score. Sliding window gene sets had a size of 50 genes, starting from the top of the filtered gene list and moved by 10 genes for each subsequent window. Each window gene set score was compared with a null distribution generated from randomly selected genes (see below). Top gene sets were determined based on a nominally significant sliding window enrichment for at least one tissue structure above the 0.95-quantile of the null in the dataset (if window scores were outside the maximum value of the null distribution, this was selected as a cut-off instead).

### Average gene expression level calculation and permutations

The relative average gene expression level score was calculated individually for each dataset first for the entire tissue slide, using dataset specific filtered gene lists for i) all genetically associated genes, ii) top genetically associated genes (see above) and iii) drug target associated genes, with the *AddModuleScore* function in Seurat. The pre-filtering of gene lists ensured that all genes used for score calculation were present and contributed to the dataset-wide relative scoring. The average expression level score was then subsetted into the spatially defined tissue structures and the median score from each such structure was extracted. We generated 10,000 null distributions by randomly sampling from all genes in the dataset without replacement using the same number of genes in each gene list. We calculated the mean expression level score for each structure with the *AddModuleScore* function. For each tissue structure (or cluster), a permutation p-value was calculated by comparing the median score, generated from dataset specific filtered gene lists for i) all genetically associated genes, ii) top genetically associated genes or iii) drug target associated genes, to comparable null distributions. We calculated the permutation p-value as:

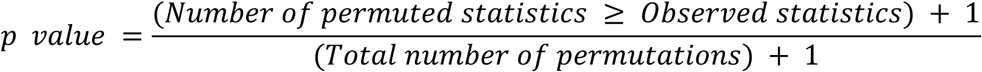

Multiple testing correction, using Bonferroni, was applied to all p-values based on the total number of tissue structures (or clusters) in a dataset.

### Downstream analyses

We used the Storey-Tibshirani method^68^ implemented in the R-package qvalue (version 2.28.0) with the *pi0est* function and pi0.method set to bootstrap^69^ to estimate the proportion of true positives (or non-null hypotheses) among sets of hypothesis tests. This was done by first estimating π_0_, the proportions of true null hypotheses, and calculating π_1_ = 1 - π_0_.

Spearman correlations were calculated in R using the *cor* function. Heatmaps were generated using the *Heatmap* function from the ComplexHeatmap^70^ (version 2.15.4) R-package, using hierarchical clustering with complete linkage to find similar clusters.

### Single-cell analyses

Single-cell deconvolution of the SRT dataset (brain_mouse_10X) was performed using the *RunNNLS* function from the Semla^50^ R-package, with a randomly downsampled (without replacement, seed 42) subset of cells from the Allen Brain Atlas of the mouse cortex and hippocampus (ref). The subsetted cells were restricted to a maximum of 250 single-cells from each subclass, with the exception of Meis2 (n=20) and VLMC (n=152) from which all available cells were included.

To reduce sparseness and increase the power of the single-cell and single-nuclei datasets, we employed a meta cell strategy. The mouse and human brain datasets were pre-processed according to the following steps and we used the same cluster annotations as the referred to publications of the respective datasets. The single-cell mouse dataset from Allen Brain Atlas of the mouse cortex and hippocampus (ref) was randomly downsampled as described above. We then applied empirical filtering of cells (200 < genes < 8000, 500 < UMI < 35000, mitochondrial-% <= 10, ribosomal-% <= 15) and removed genes with less than 100 UMI counts. The single-nuclei adult human brain datasets^49^ for AllNeurons and AllNonNeurons were randomly downsampled (without replacement, seed 123) to a maximum of 250 single-nucleus from each supercluster where subsetted cells were restricted to a maximum of 250 single-cells from each supercluster. We then merged the subsetted datasets for AllNeurons and AllNonNeurons and applied empirical filtering of cells (200 < genes < 15000, 500 < UMI < 125000, mitochondrial-% <= 5, ribosomal-% <= 4) and removed genes with less than 100 UMI counts.

Before generating meta cells, we removed cell types with less than 50 cells (i.e., Meis2). Datasets were then split based on cell type annotation and individually analyzed in Seurat by normalization (*SCTransform*), identification of variable genes (*FindVariableFeatures*) and principal component analysis (*RunPCA*). Clustering was performed using the *FindNeighbors* function and the top 10 principal components. Then using the *FindClusters* function starting at a resolution of 0.1 and iterating through an increase of 0.1 in resolution until yielding a mean number of cells per cluster of ∼10. Each cluster within a cell type was then aggregated into a meta cell, and subsequently all meta cells were merged back into one dataset. We used the same cluster annotations as the publications of the respective datasets.

We applied STEAM to calculate gene set enrichment for ALZ, ADHD, ASD, BIP and SCZ. Here, using the sliding window approach to identify top genes and reduce the signal from non-coexpressing genes. Enrichments across ranked gene lists were tested for multiple correction based on number of windows and cell types, using Benjamini & Hochberg. In addition, we performed a complementary enrichment analysis, testing for enrichment of genetically associated genes among the upregulated differentially expressed genes for each cell type using a one-sided Fisher’s exact test with function *fisher.test* and alternative set to greater. The complementary enrichment analysis was performed using all and top genes identified with STEAM.

### Spatial drug repurposing candidates

We utilized the drug target gene lists from the Open Targets database^7,8^, and computed Spearman correlations to look for disease-associated genes grouping in spatially colocalized subsets, which we previously identified as significantly enriched based on the genetically associated gene lists. We included: all genetically associated genes used to determine the trait-specific enrichment, and all drug target genes from the 32 complex traits within this study that also had a genetic association to the trait investigated. We labeled the genes according to their association with the trait investigated, as i) genetic, ii) a known drug target, iii) a “brain trait” or iv) a “non-brain trait” as depicted in Figure 1a. We constructed correlation-based heatmaps and searched for clusters containing genes with spatially defined expression profiles that implicated the same tissue structures we previously found to have trait-specific enrichments (Supplementary Table 5). This enabled us to select spatially informed drug repurposing candidates from the heatmap clusters and check which other traits these drug repurposing candidates were associated with (Supplementary Table 8).

## Data availability

Links and metadata to published spatial transcriptomics datasets used in this study can be found in Supplementary Table 1. The 10X Genomics dataset for mouse cortex can be found at https://www.10xgenomics.com/datasets/mouse-brain-section-coronal-1-standard-1-1-0 and for mouse kidney at https://www.10xgenomics.com/datasets/mouse-kidney-section-coronal-1-standard-1-1-0. The scRNA-seq Allen Brain Atlas of the mouse cortex and hippocampus^48^, can be found at https://portal.brain-map.org/atlases-and-data/rnaseq/mouse-whole-cortex-and-hippocampus-10x. The snRNA-seq of the adult human brain^49^, can be found at https://cellxgene.cziscience.com/collections/283d65eb-dd53-496d-adb7-7570c7caa443.

## Code availability

The STEAM algorithm’s computational workflow (Supplementary Fig. 30) and code can be found on Github (https://github.com/kvastad/STEAM), and the STEAM R-package will be available upon the publication of the manuscript.

## Acknowledgements

We thank Enikő Lázár, Ludvig Larsson, Javier Escudero Morlanes and Lovisa Franzén for valuable discussions. This work was supported by a grant from the Knut and Alice Wallenberg Foundation to SciLifeLab for research in Data-driven Life Science, DDLS (grants: KAW 2020.0239 and KAW 2017.0003), and by the National Bioinformatics Infrastructure Sweden (NBIS) at SciLifeLab. The computations were enabled by resources in project (SNIC 2022-5-560 and NAISS 2023-5-517) provided by the National Academic Infrastructure for Supercomputing in Sweden (NAISS) at UPPMAX, funded by the Swedish Research Council through grant agreement no. 2022-06725.

## Author contribution statement

LK and TL designed the study and drafted the manuscript. LK and AK developed the methodology and code. LK performed the core data analysis. LK, AK, CC, and TC performed initial data analysis. R.S. and L.K. co-designed the R-package. R.S. wrote the R-package. All authors reviewed, read, and approved the final manuscript.

## Ethics declarations

T.L. is an advisor and has equity in Variant Bio, was a paid advisor to GSK and has received speaker honoraria from Abbvie and Merck.

