## Supplementary Information for "Spatial transcriptomics and genetically implicated genes identify putative causal tissue structures for complex traits"

* Current affiliation
Department of Cell and Molecular Biology, Karolinska Institute, Solna, Sweden

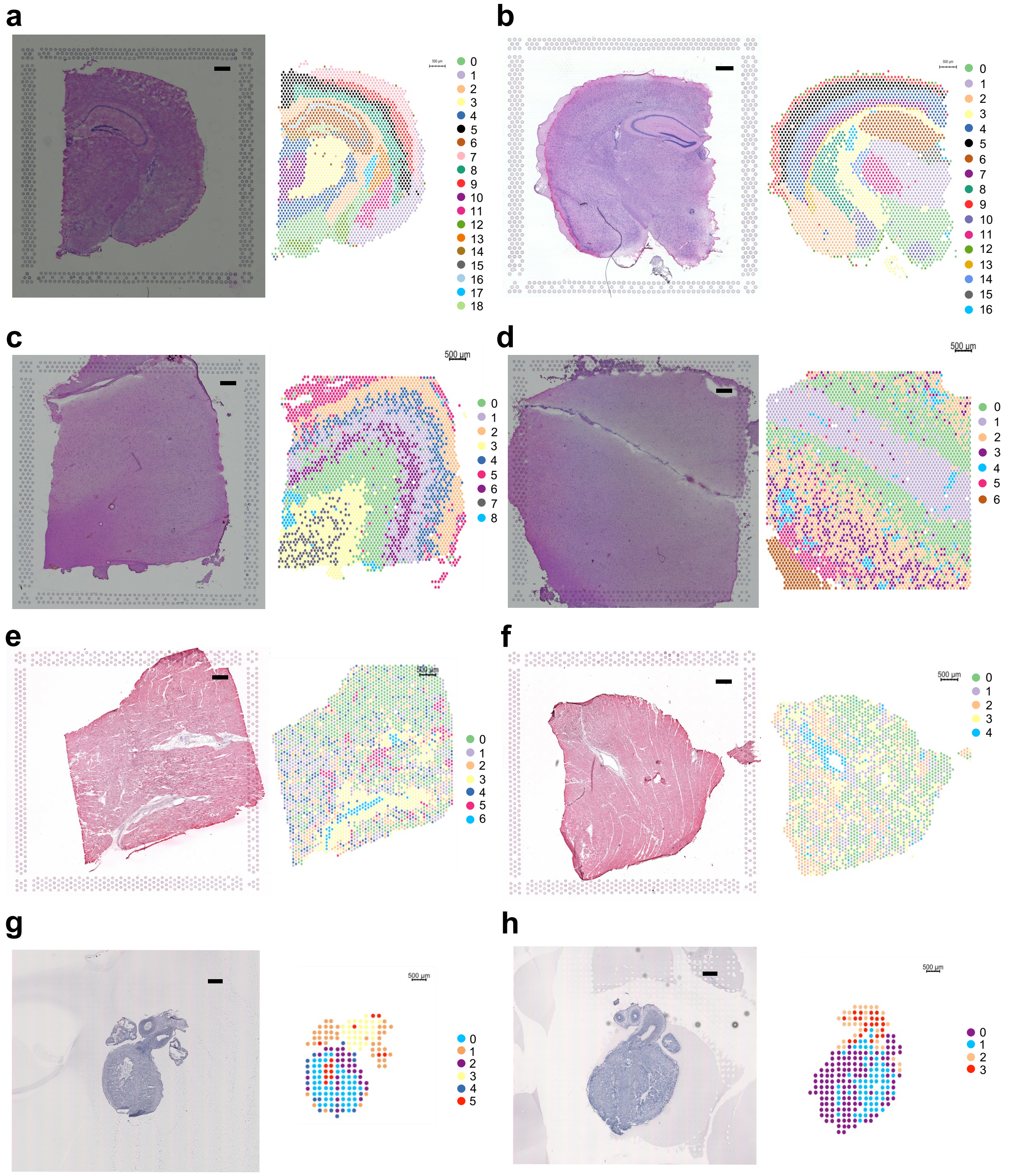


**Supplementary Figure 1. HE and spatial structures of brain and heart.** Dataset **a)** brain_mouse_10X, **b)** brain_mouse_cell2location, **c)** brain_human_151673, **d)** brain_human_151509, **e)** heart_human_adult_ACH003, **f)** heart_human_adult_ACH004, **g)** heart_human_pcw6.5_nr8, **h)** heart_human_pcw9_nr16. Scale bar 500µm.

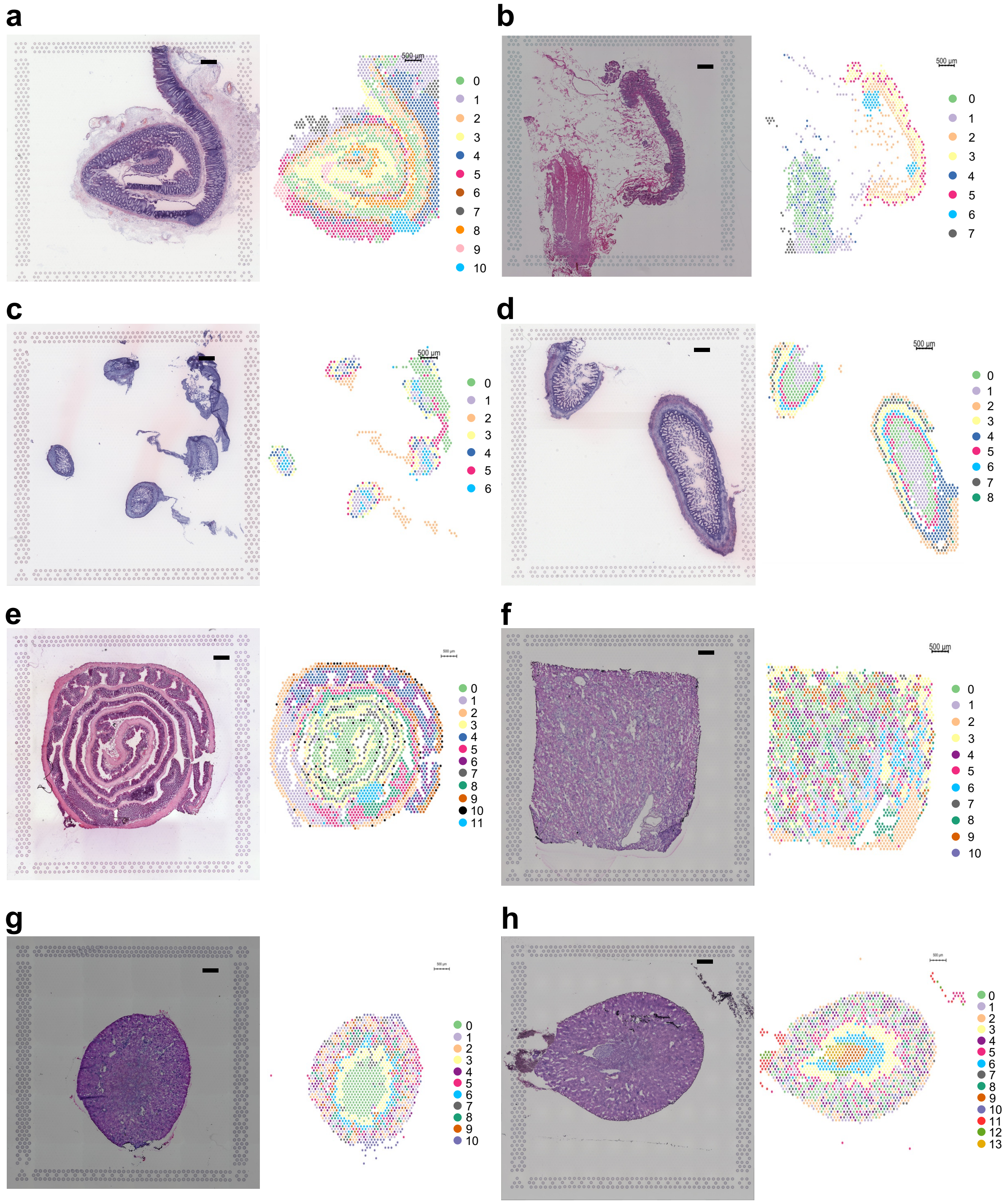


**Supplementary Figure 2. HE and spatial structures of intestine and kidney. a)** intestine_human_adult_HFGA_A1, **b)** intestine_human_adult_RRST_349_B1, **c)** intestine_human_pcw12_HFGA_A8, **d)** intestine_human_pcw19_HFGA_A4, **e)** intestine_mouse_healing_ctrl, **f)** kidney_human_JCI, **g)** kidney_mouse_10X, **h)** kidney_mouse_JCI. Scale bar 500μm.

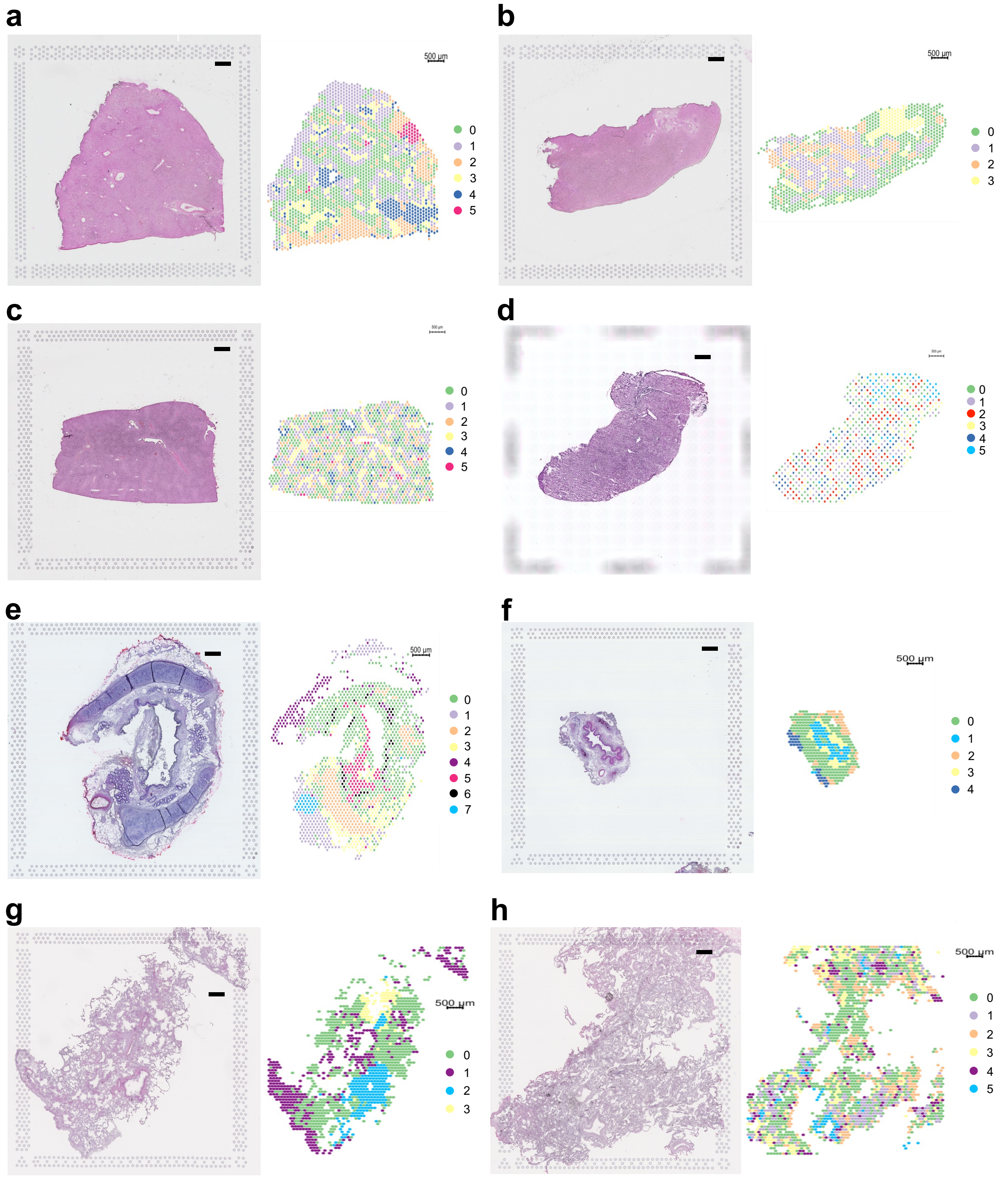


**Supplementary Figure 3. HE and spatial structures of liver and lung. a)** liver_human_JBO0018, **b)** liver_human_JBO0022, **c)** liver_mouse_JBO002, **d)** liver_mouse_zonation_C1, **e)** lung_bronchi_human_A42, **f)** lung_bronchi_human_A50, **g)** lung_parenchyma_human_A37, **h)** lung_parenchyma_human_A48. Scale bar 500µm.

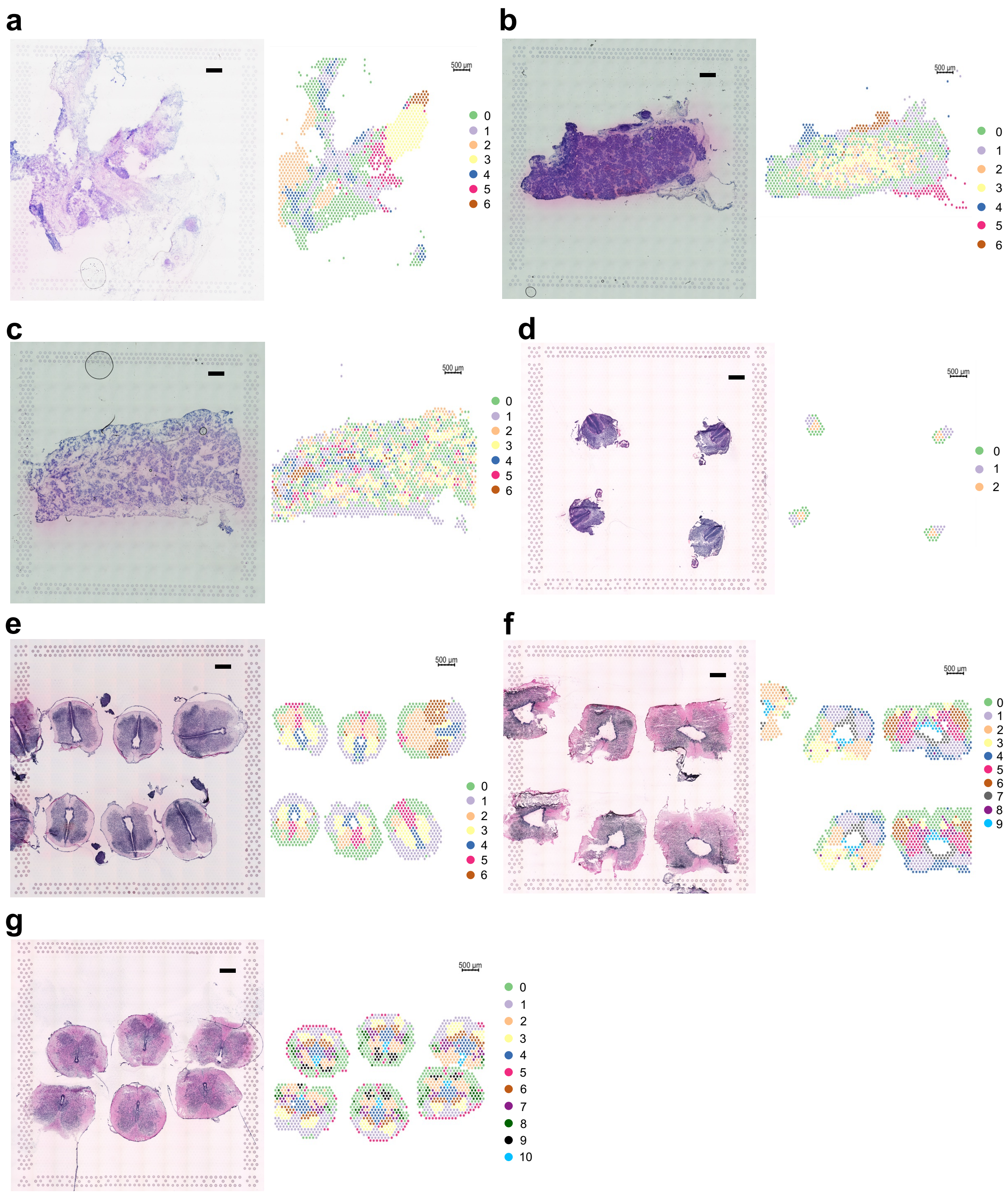


**Supplementary Figure 4. HE and spatial structures of pancreas and spinal cord. a)** pancreas_human_pcw15_S1, **b)** pancreas_human_pcw18_S1, **c)** pancreas_human_pcw20_S1, **d)** spinal_cord_human_pcw5_302B1, **e)** spinal_cord_human_pcw8_107B1, **f)** spinal_cord_human_pcw9_288A1, **g)** spinal_cord_human_pcw12_290D1. Scale bar 500µm.

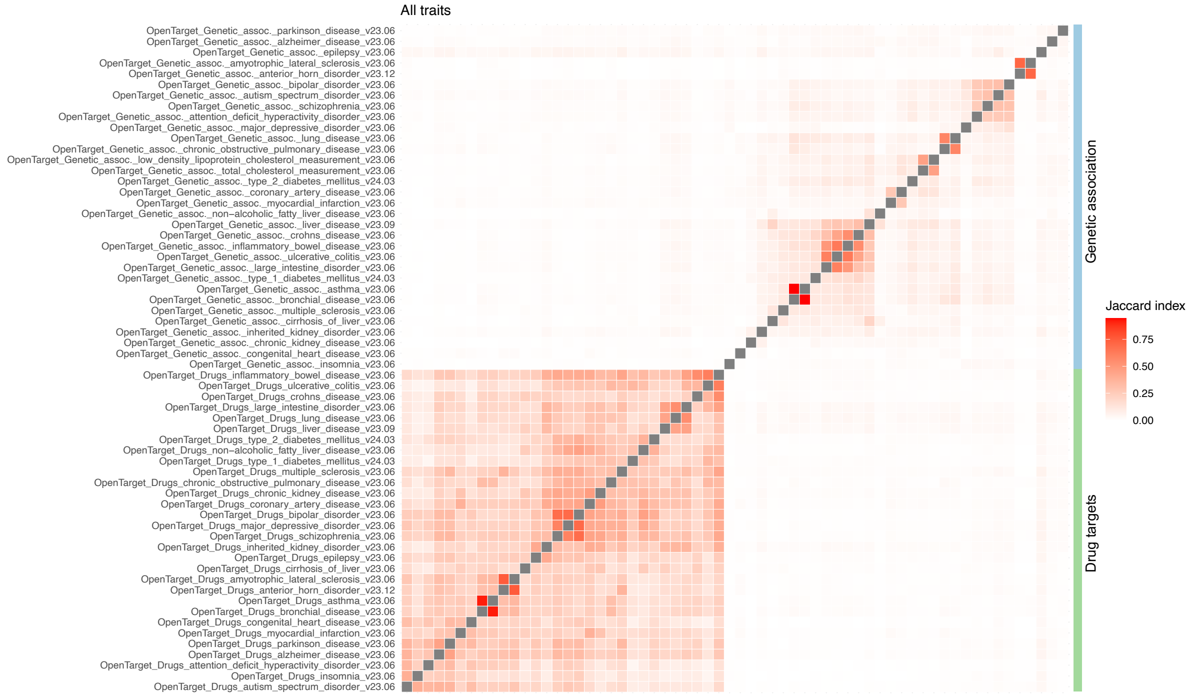


**Supplementary Figure 5. Jaccard index between all gene lists.** Open targets gene lists for genetically associated genes and drug targets for each trait. Open targets release version is indicated by version-year-month.

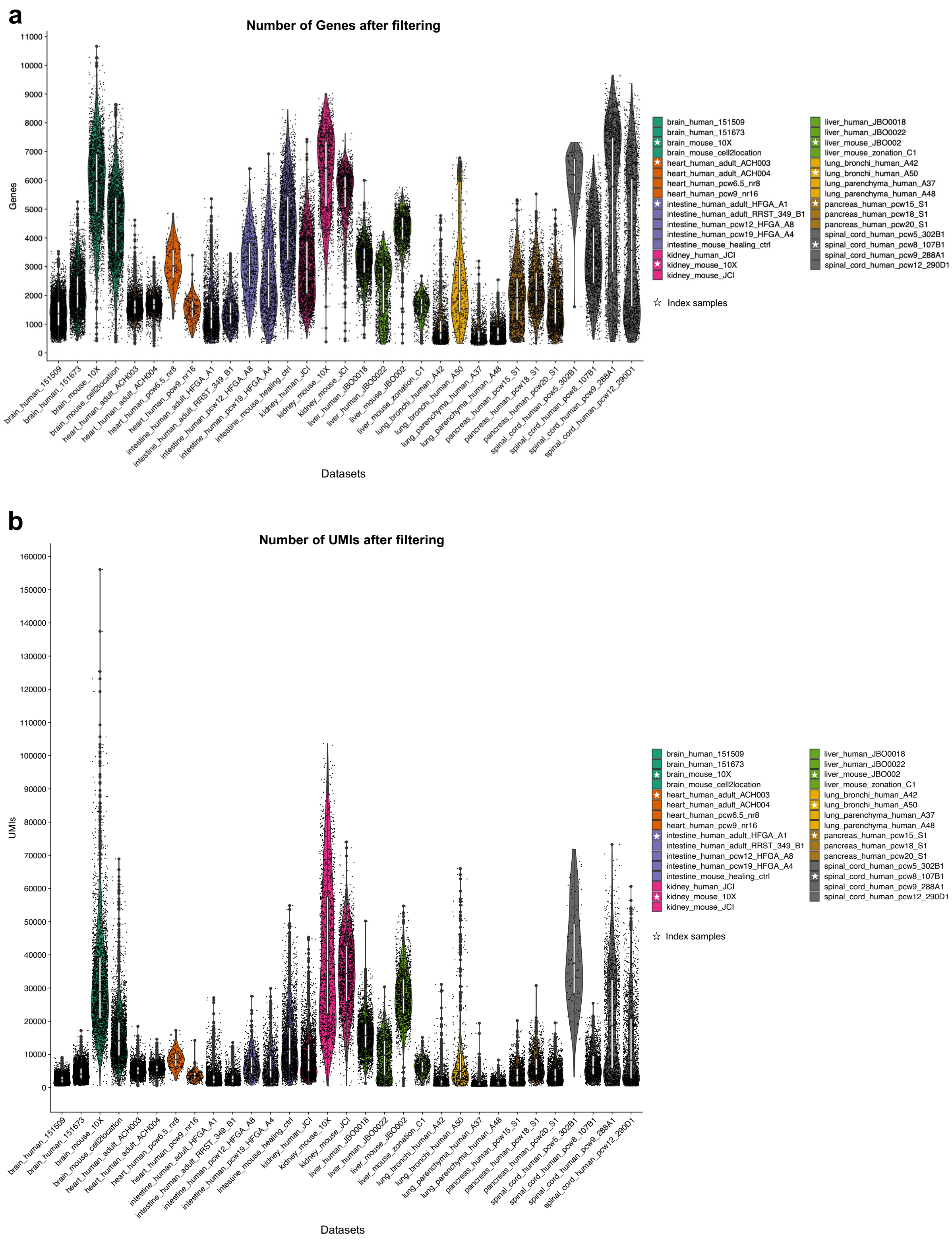


**Supplementary Figure 6. Data distribution across datasets after quality filtering.** Index samples are indicated by a star in the legend, **a)** number of genes and **b)** number of UMIs.

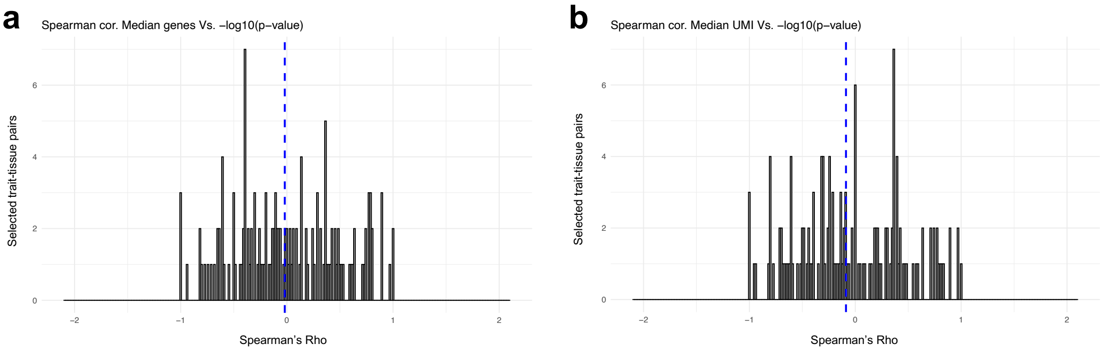


**Supplementary Figure 7. Spearman correlations across selected trait-tissue pairs for genetically associated genes.** Spearman correlations were calculated across spatial structures between nominal p-values and **a)** genes or **b)** UMIs. This was done for each selected trait-tissue pair in each spatial dataset, as depicted in Figure 1a. Blue dotted line, median.

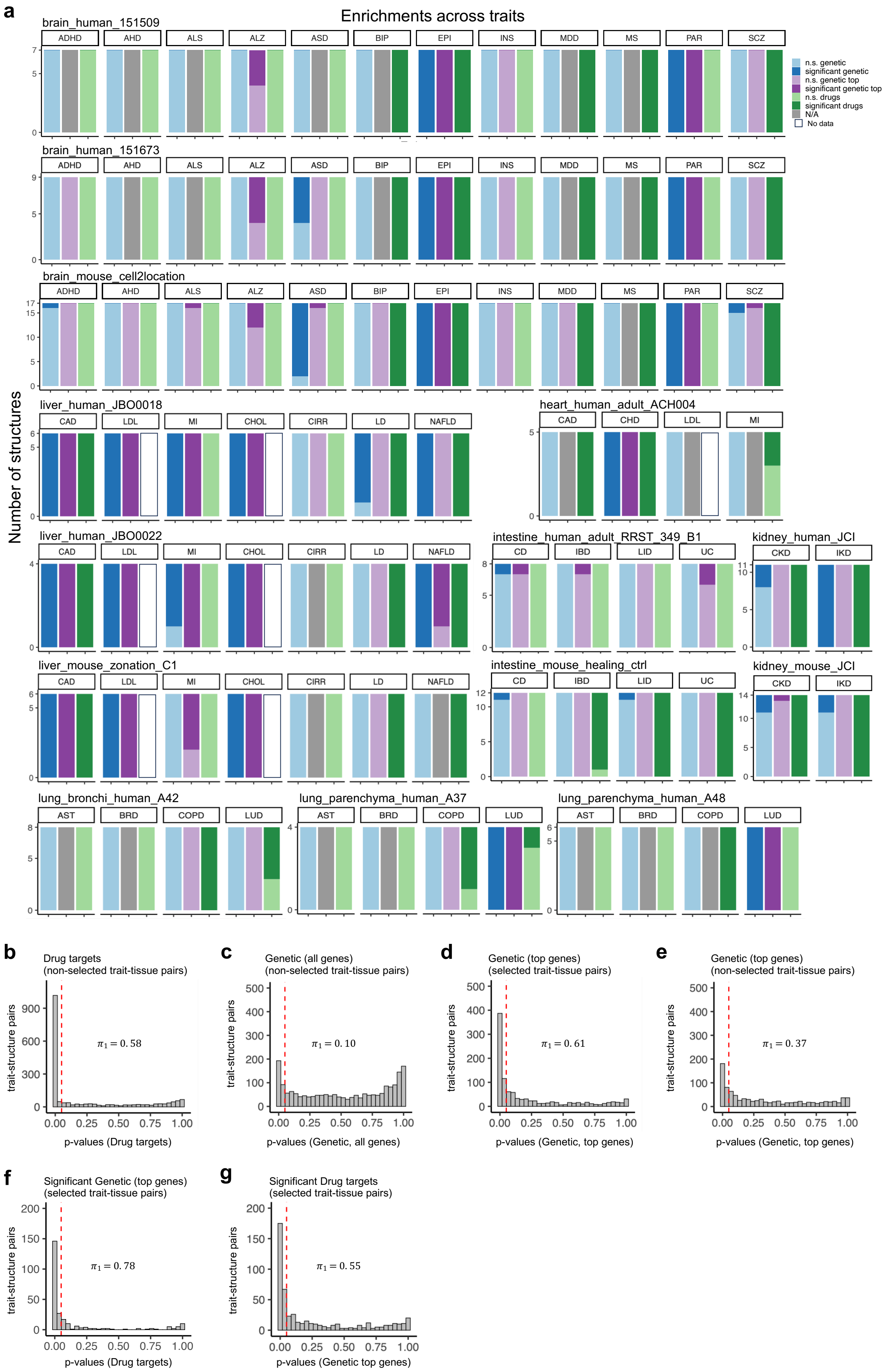
a
Enrichments across traits
Number of structures
brain_human_151509
brain_human_151673
brain_mouse_cell2location
liver_human_JBO0018
heart_human_adult_ACH004
liver_human_JBO0022
intestine_human_adult_RRST_349_B1
kidney_human_JCI
liver_mouse_zonation_C1
intestine_mouse_healing_ctrl
kidney_mouse_JCI
lung_bronchi_human_A42
lung_parenchyma_human_A37
lung_parenchyma_human_A48
ADHD AHD ALS ALZ ASD BIP EPI INS MDD MS PAR SCZ
CAD LDL MI CHOL CIRR LD NAFLD
CAD CHD LDL MI
CD IBD LID UC
CKD IKD
AST BRD COPD LUD
n.s. genetic
significant genetic
n.s. genetic top
significant genetic top
n.s. drugs
significant drugs
N/A
No data
b
Drug targets
(non-selected trait-tissue pairs)
$\pi_1 = 0.58$
p-values (Drug targets)
c
Genetic (all genes)
(non-selected trait-tissue pairs)
$\pi_1 = 0.10$
p-values (Genetic, all genes)
d
Genetic (top genes)
(selected trait-tissue pairs)
$\pi_1 = 0.61$
p-values (Genetic, top genes)
e
Genetic (top genes)
(non-selected trait-tissue pairs)
$\pi_1 = 0.37$
p-values (Genetic, top genes)
f
Significant Genetic (top genes)
(selected trait-tissue pairs)
$\pi_1 = 0.78$
p-values (Drug targets)
g
Significant Drug targets
(selected trait-tissue pairs)
$\pi_1 = 0.55$
p-values (Genetic top genes)
trait-structure pairs

**Supplementary Figure 8. Putative causal tissue structures. a)** Enrichments across traits in all adult non-index samples for each tissue. **b-g)** Enrichments of trait-structure pairs measured as $\pi_1$ statistics using nominal p-values (red dotted line = 0.05). **b)** Non-selected trait-tissue pairs for drug targets. **c)** Non-selected trait-tissue pairs for all genetically implicated genes. **d)** Selected and **e)** non-selected trait-tissue pairs for top genes. **f)** Drug target implicated genes for structures with significantly enriched top genes for the same trait and **g)** top genetically implicated genes for structures with significantly enriched drug targets for the same trait.

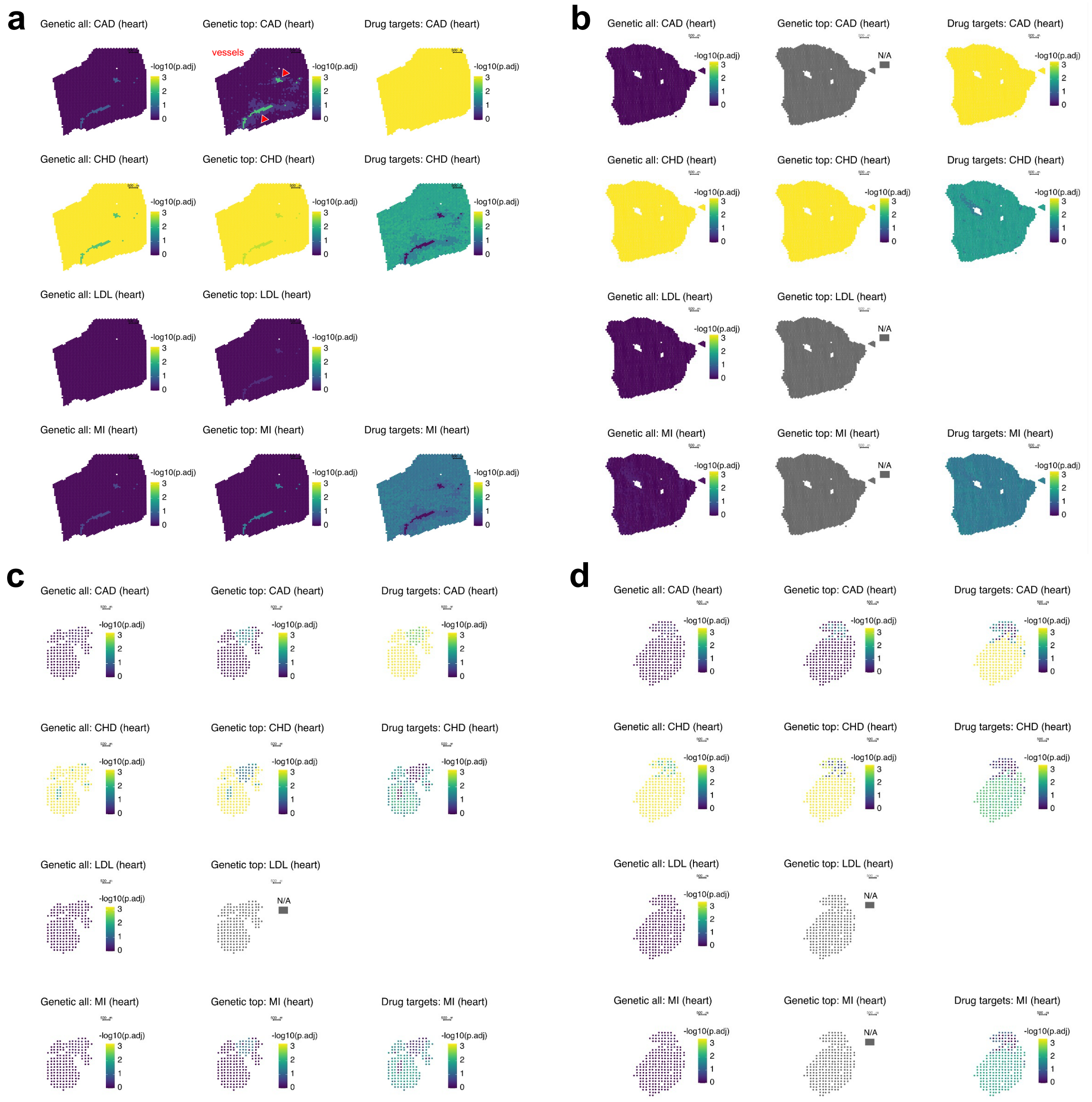


**Supplementary Figure 9. Spatial trait enrichments in the human developmental and adult heart.** Visualized with trait-specific Bonferroni adjusted permutation p-values in datasets **a)** heart_human_adult_ACH003, with arrows (red) indicating spatial structure 6 for vessels for CAD. **b)** heart_human_adult_ACH004, **c)** heart_human_pcw6.5_nr8, **d)** heart_human_pcw9_nr16. Scale bar 500µm.

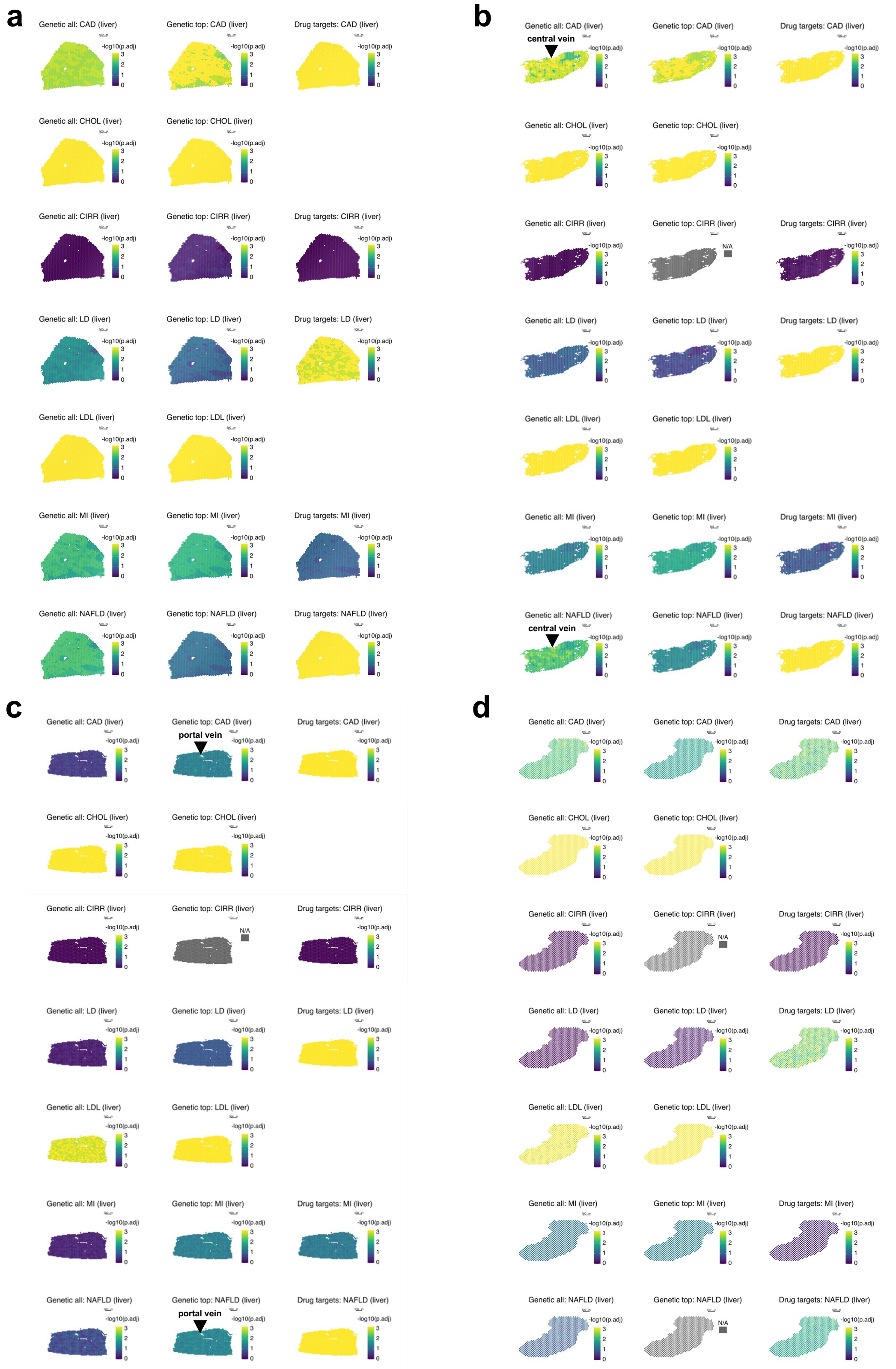


**Supplementary Figure 10. Spatial trait enrichments in the human and mouse liver.** Visualized with trait-specific Bonferroni adjusted permutation p-values in datasets **a)** liver_human_JBO018, **b)** liver_human_JBO022 with arrow indicating one central vein from spatial structure 2 for NAFLD, **c)** liver_mouse_JBO002 with arrow indicating one portal vein from spatial structure 4 for NAFLD, **d)** liver_mouse_zonation_C1. Scale bar 500µm.

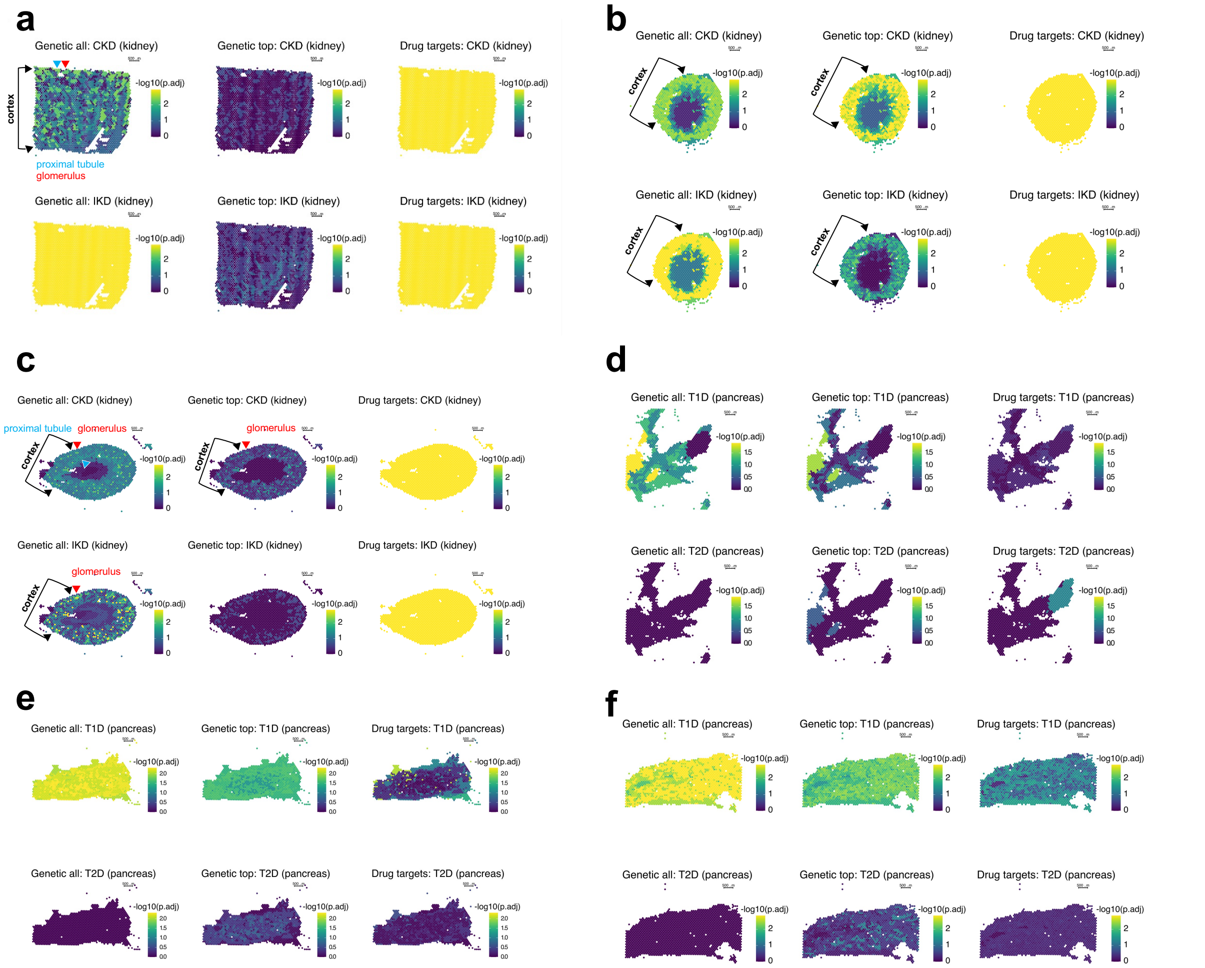


**Supplementary Figure 11. Spatial trait enrichments in the human and mouse kidney and the human developmental pancreas.** Visualized with trait-specific Bonferroni adjusted permutation p-values in datasets **a)** kidney_human_JCI with arrow (blue) indicating spatial structure 3 as proximal tubule and arrow (red) spatial structure 4 as glomerulus for CKD, **b)** kidney_mouse_10X with arrows indicating the larger cortex structure, **c)** kidney_mouse_JCI with arrows (black) indicating the larger cortex structure, with arrow (blue) indicating spatial structure 1 and 5 as proximal tubule and with arrows (red) spatial structure 8 as glomerulus, **d)** pancreas_human_pcw15_S1, **e)** pancreas_human_pcw18_S1, **f)** pancreas_human_pcw20_S1. Scale bar 500μm.

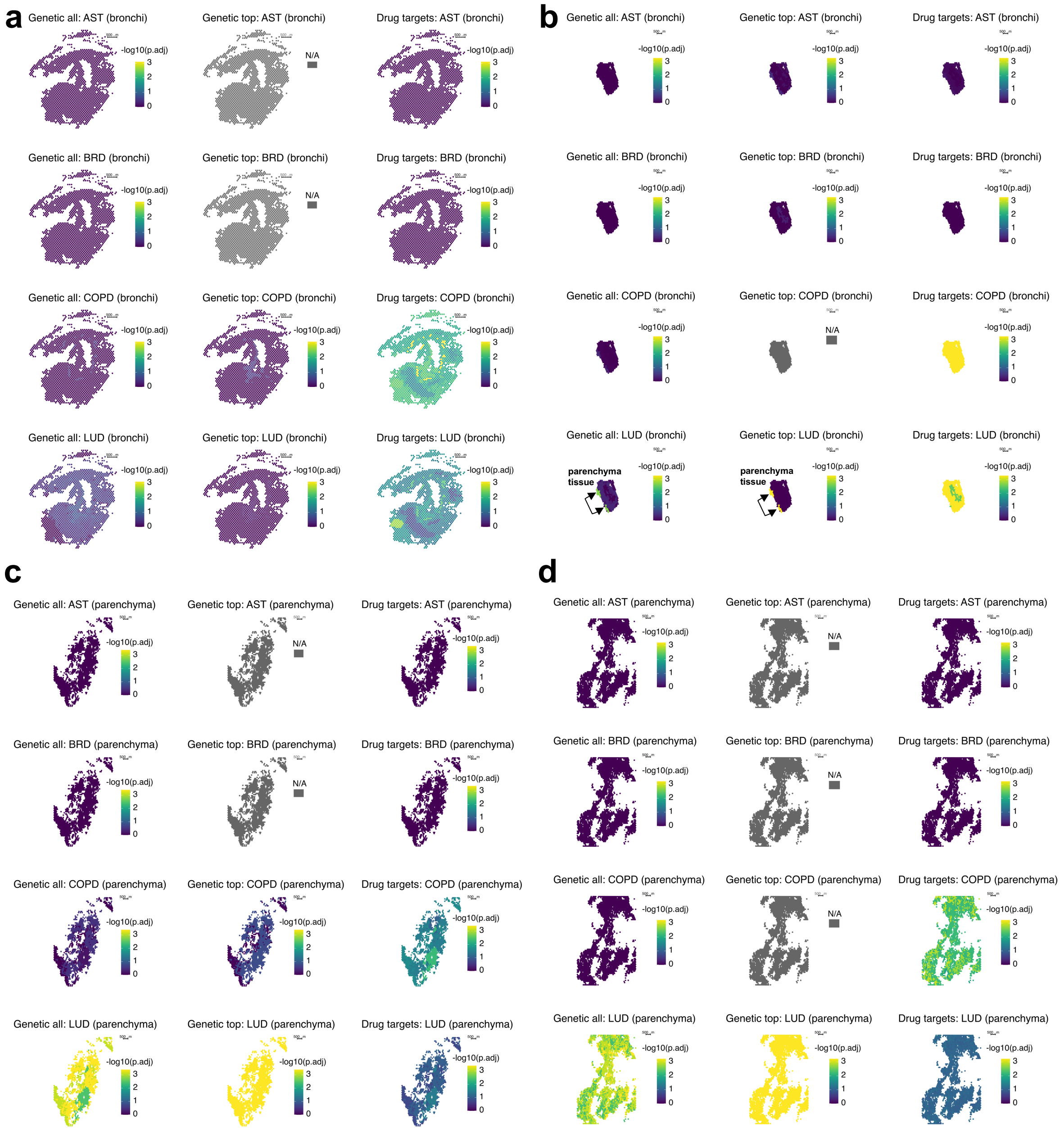


**Supplementary Figure 12. Spatial trait enrichments in the human lung.** Visualized with trait-specific Bonferroni adjusted permutation p-values in datasets **a)** lung_bronchi_human_A42, **b)** lung_bronchi_human_A50 with arrows indicating spatial structure 4 for parenchyma tissue for LUD, **c)** lung_parenchyma_human_A37, **d)** lung_parenchyma_human_A48. Scale bar 500µm.

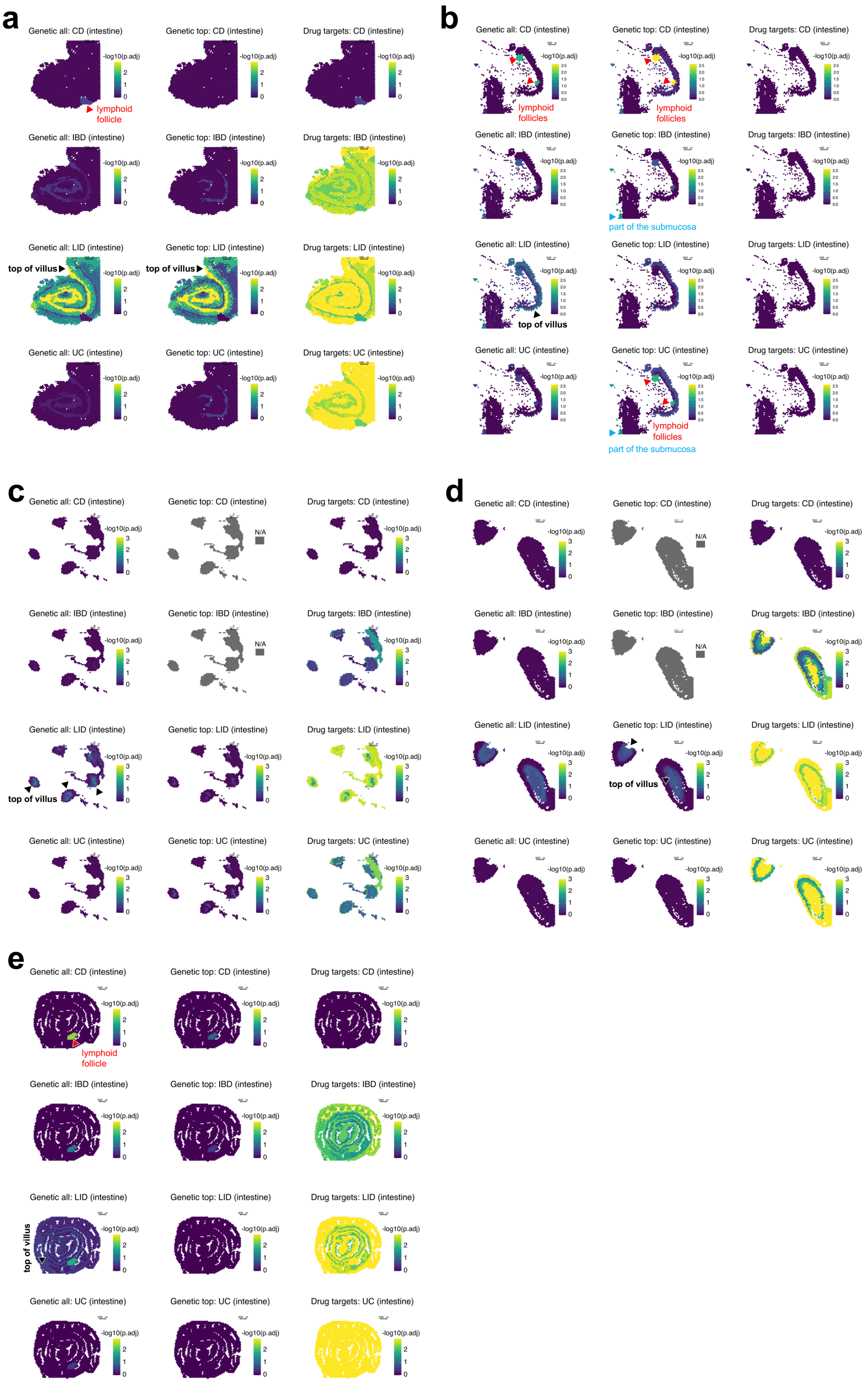


**Supplementary Figure 13. Spatial trait enrichments in the human and mouse intestine.** Visualized with trait-specific Bonferroni adjusted permutation p-values in datasets **a)** intestine_human_adult_HFGA_A1 with arrows (black) indicating spatial structure 3 and 8 for top of the villus for LID and arrow (red) indicating spatial structure 10 for lymphoid follicle for CD. **b)** intestine_human_adult_RRST_349_B1 with arrows (red) indicating spatial structure 6 for lymphoid follicles for CD and UC, and arrow (blue) indicating spatial structure 7 for part of the submucosa for IBD and UC, and arrow (black) indicating spatial structure 5 for top of the villus for LID. **c)**

intestine_pcw12_HFGA_A8 with arrows (black) indicating spatial structure 6 for top of the villus for LID. **d)** intestine_pcw19_HFGA_A4 with arrows (black) indicating spatial structure 1 for top of the villus for LID. **e)** intestine_mouse_healing_ctrl with arrow (black) indicating spatial structure 7 for top of the villus for LID and arrows (red) indicating spatial structure 11 for lymphoid follicle for CD. Scale bar 500μm.

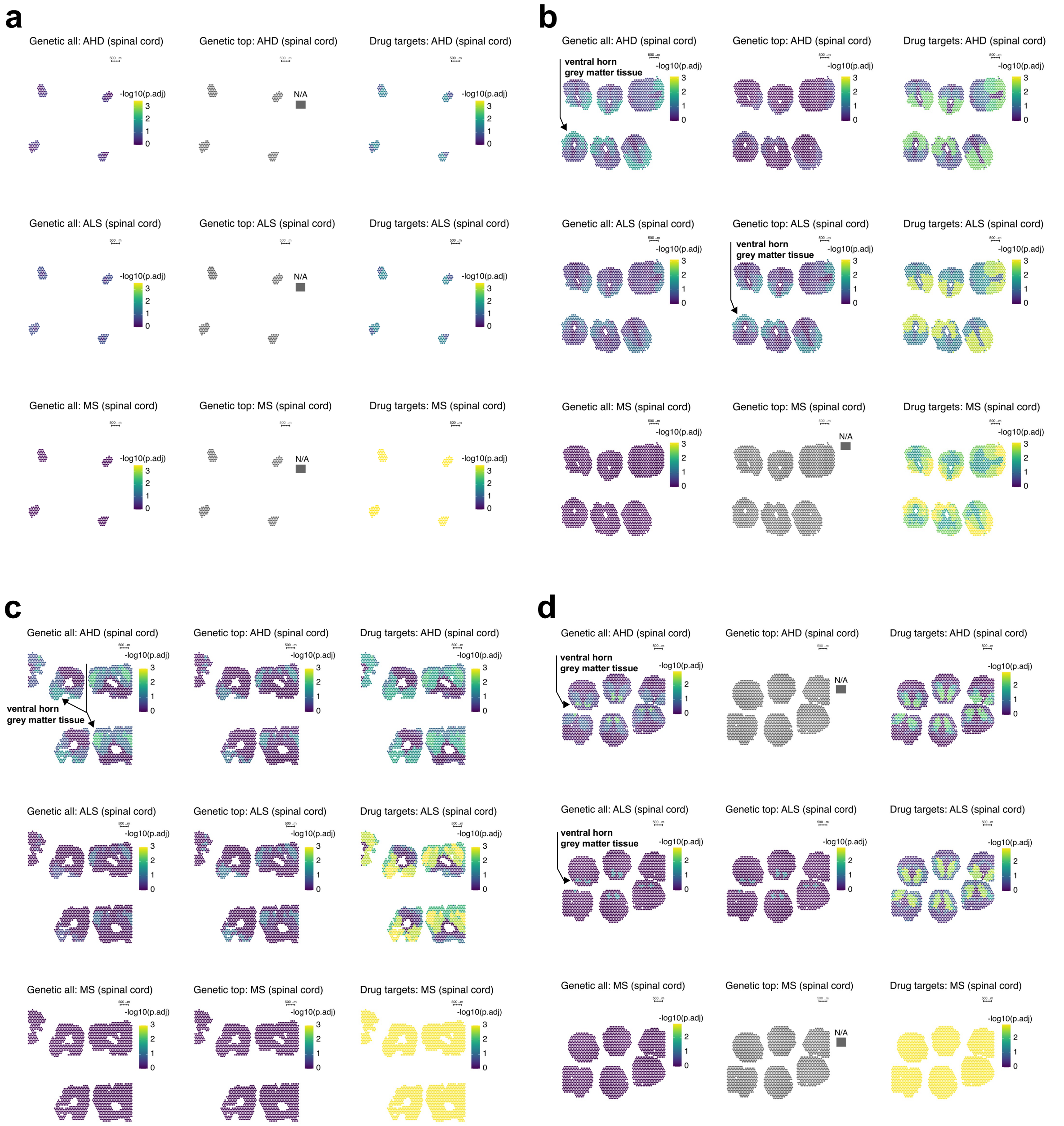


**Supplementary Figure 14. Spatial trait enrichments in the human developmental spinal cord.** Visualized with trait-specific Bonferroni adjusted permutation p-values in datasets **a)** spinal_cord_human_pcw5_302B1, **b)** spinal_cord_human_pcw8_107B1 with arrows indicating spatial structure 1 for the ventral horn grey matter tissue in one of the sections for AHD and ALS, **c)** spinal_cord_human_pcw9_288A1 with arrow indicating spatial structures 6 and 3 for the ventral horn grey matter tissue in one of the sections for AHD, **d)** spinal_cord_human_pcw12_290D1 with arrows indicating spatial structure 9 for the ventral horn grey matter tissue in one of the sections for AHD and ALS. Scale bar 500μm.

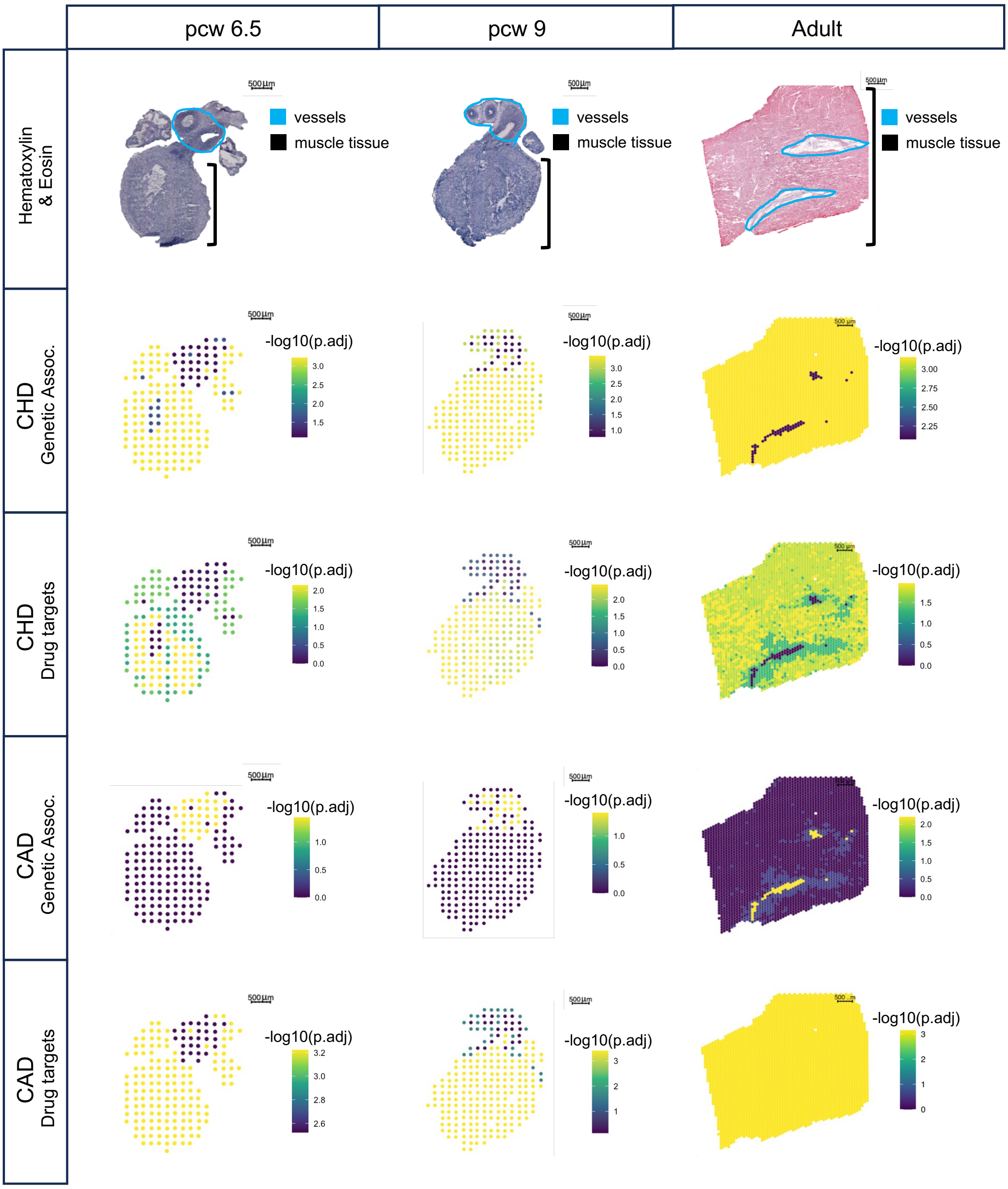


**Supplementary Figure 15. Trait-specific enrichment in spatial tissue structures of the human heart from development to adulthood.** Enrichments across traits for the heart index sample and the developmental stages using genetic top gene lists (selected based on sliding window approach) and drug targets gene list (all genes). CAD=Coronary artery disease. CHD=Congenital heart disease. pcw=post-conceptual weeks. Scale bar 500µm.

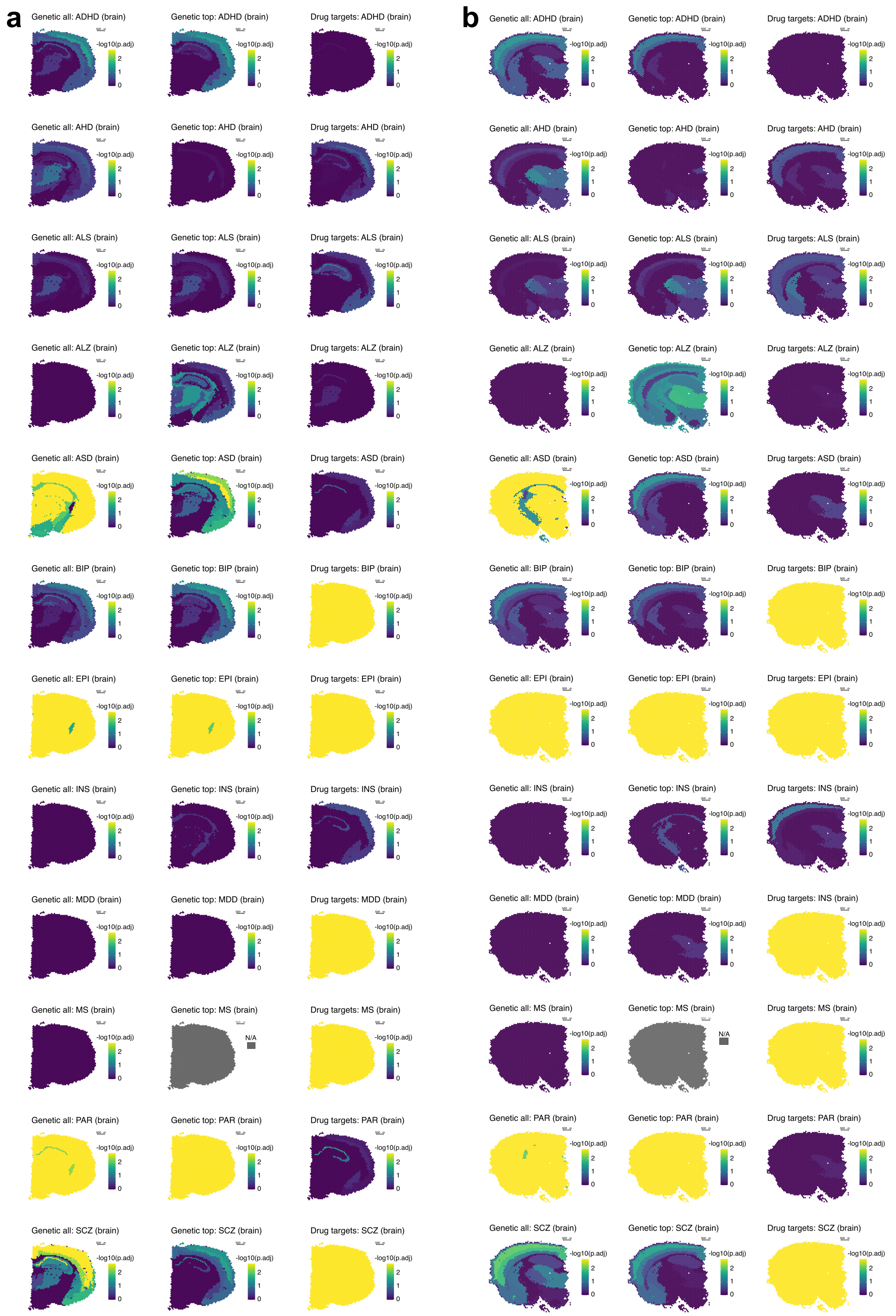


**Supplementary Figure 16. Spatial trait enrichments in the mouse brain.** Visualized with trait-specific Bonferroni adjusted permutation p-values in datasets **a)** brain_mouse_10X and **b)** brain_mouse_cell2location. Scale bar 500µm.

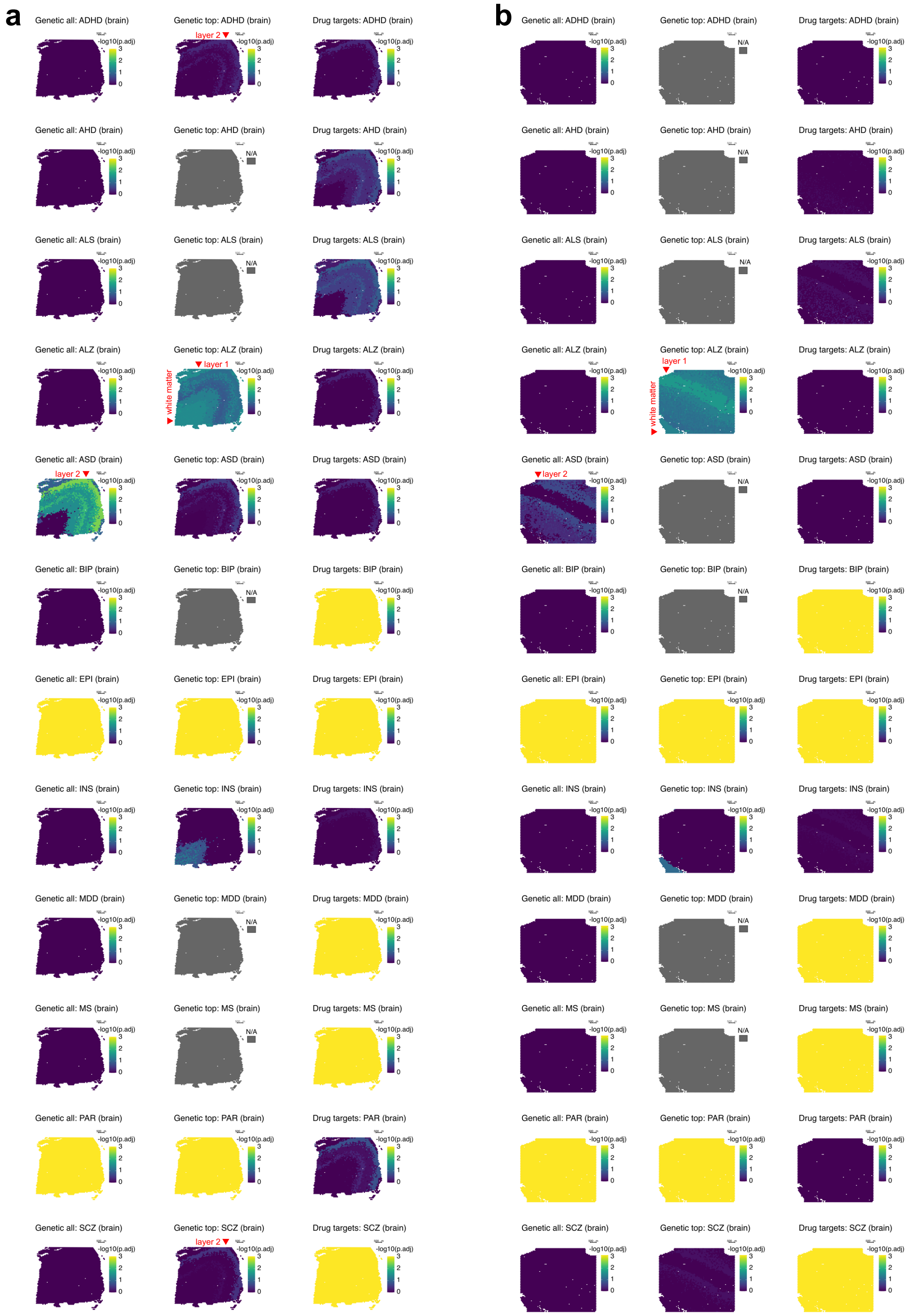


**Supplementary Figure 17. Spatial trait enrichments in the human brain.** Visualized with trait-specific Bonferroni adjusted permutation p-values in datasets **a)** brain_human_151673 with arrow indicating spatial structures 3 and 7 for white matter and spatial structure 5 for layer 1 for ALZ and spatial structures 2 for layer 2 in ADHD, ASD and SCZ and **b)** brain_human_151509 with arrow indicating spatial structure 6 for white matter and spatial structure 1 for layer 1 for ALZ and spatial structures 0 for layer 2 in ASD. Scale bar 500µm.

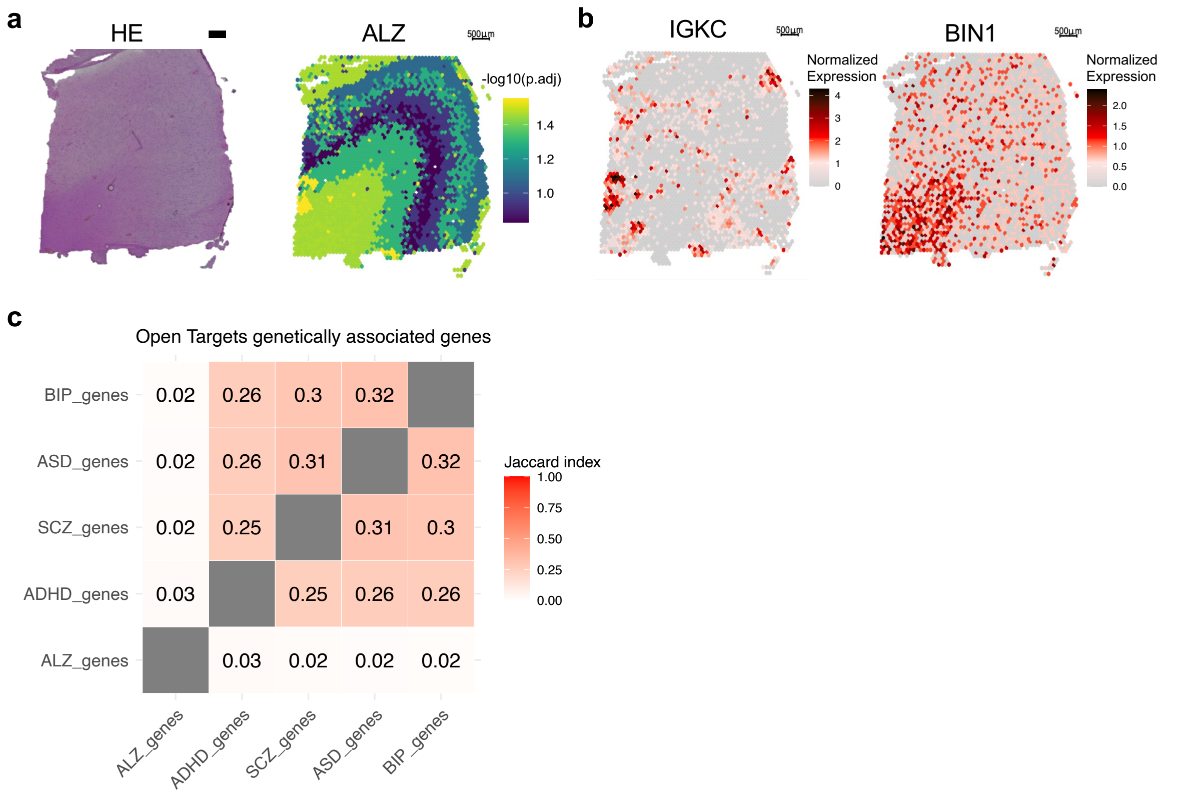


**Supplementary Figure 18. ALZ enriched structures and gene expression in the human dorsolateral prefrontal cortex. a)** HE staining (left) and spatial trait enrichment from top 60 genetically implicated genes in the human brain visualized with trait-specific Bonferroni adjusted permutation p-values in dataset brain_human_151673 (right). **b)** Normalize gene expression in the dataset brain_human_151673 marking areas with B-cell presence (left) and white matter (right). Scale bar 500µm. **c)** Overlap of trait-specific genetically associated genes from Open Targets.

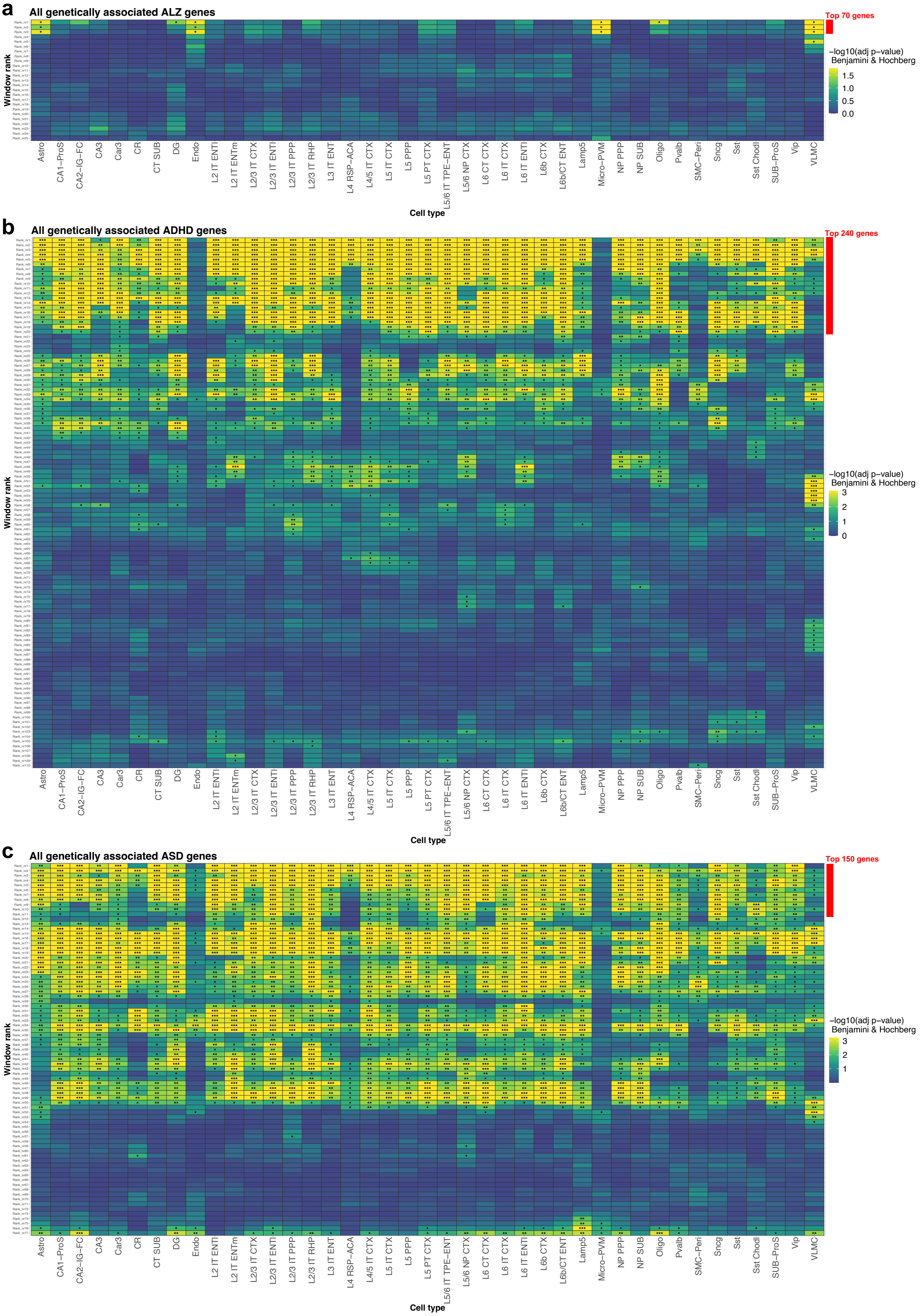


**Supplementary Figure 19. Enrichment of ALZ, ADHD and ASD across ranked gene lists in the mouse cortex and hippocampus.** Trait enrichment with STEAM in cell types (using meta cells) for genetically associated ranked gene lists across subclasses in the mouse cortex and hippocampus for **a)** ALZ, **b)** ADHD and **c)** ASD. Each ranked widow consists of 50 genes, moving by

10 genes. Multiple testing correction with Benjamini & Hochberg, * $p < 0.05$, ** $p < 0.01$ and *** $p < 0.001$. Top genes selected with STEAM marked in red. List of mouse cell subclass abbreviations can be found in Supplementary Table 6.

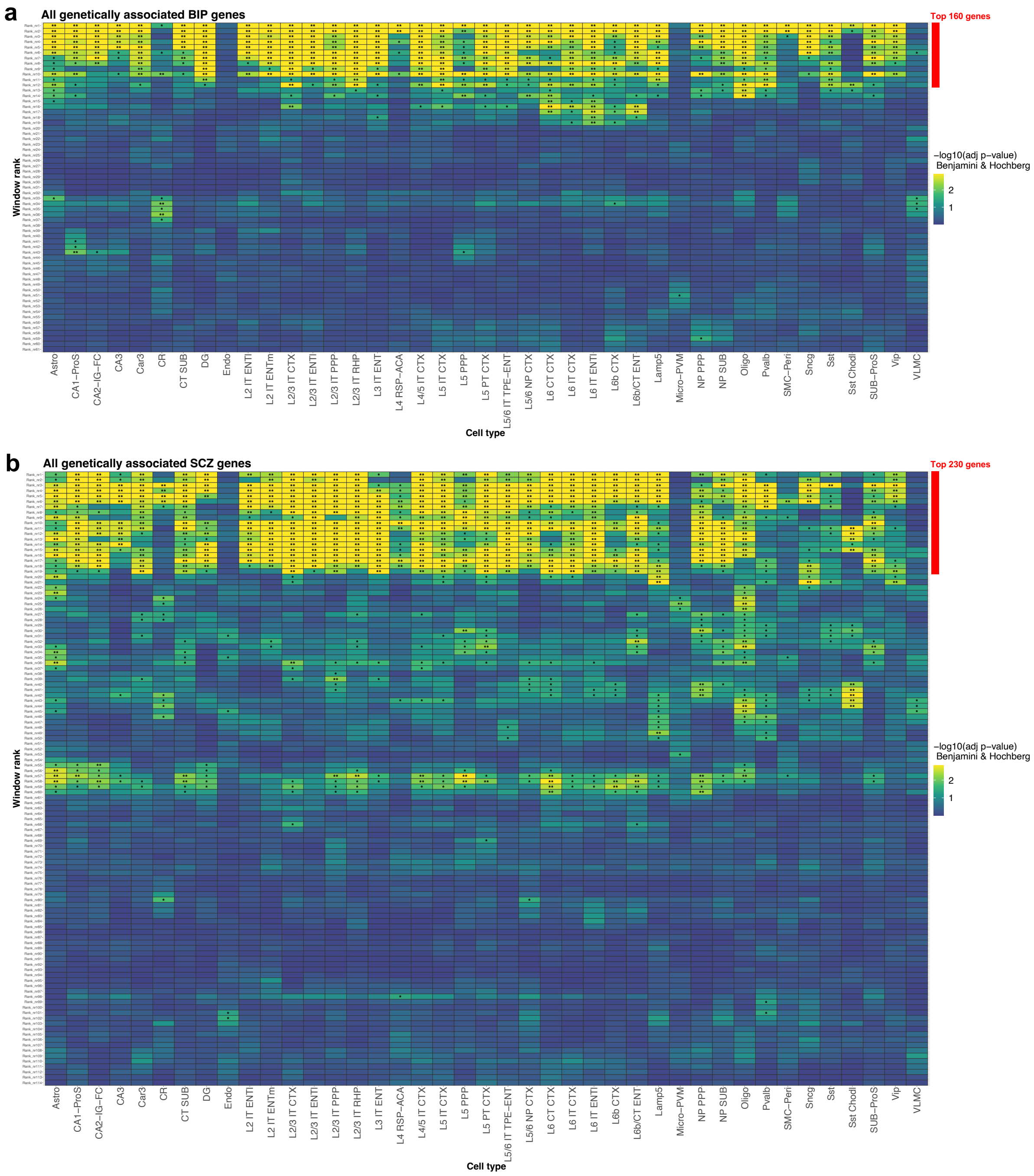


**Supplementary Figure 20. Enrichment of BIP and SCZ across ranked gene lists in the mouse cortex and hippocampus.** Trait enrichment with STEAM in cell types (using meta cells) for genetically associated ranked gene lists across subclasses in the mouse cortex and hippocampus for **a)** BIP and **b)** SCZ. Each ranked widow consists of 50 genes, moving by 10 genes. Multiple testing correction with Benjamini & Hochberg, * $p < 0.05$, ** $p < 0.01$ and *** $p < 0.001$. Top genes selected with STEAM marked in red. List of mouse cell subclass abbreviations can be found in Supplementary Table 6.

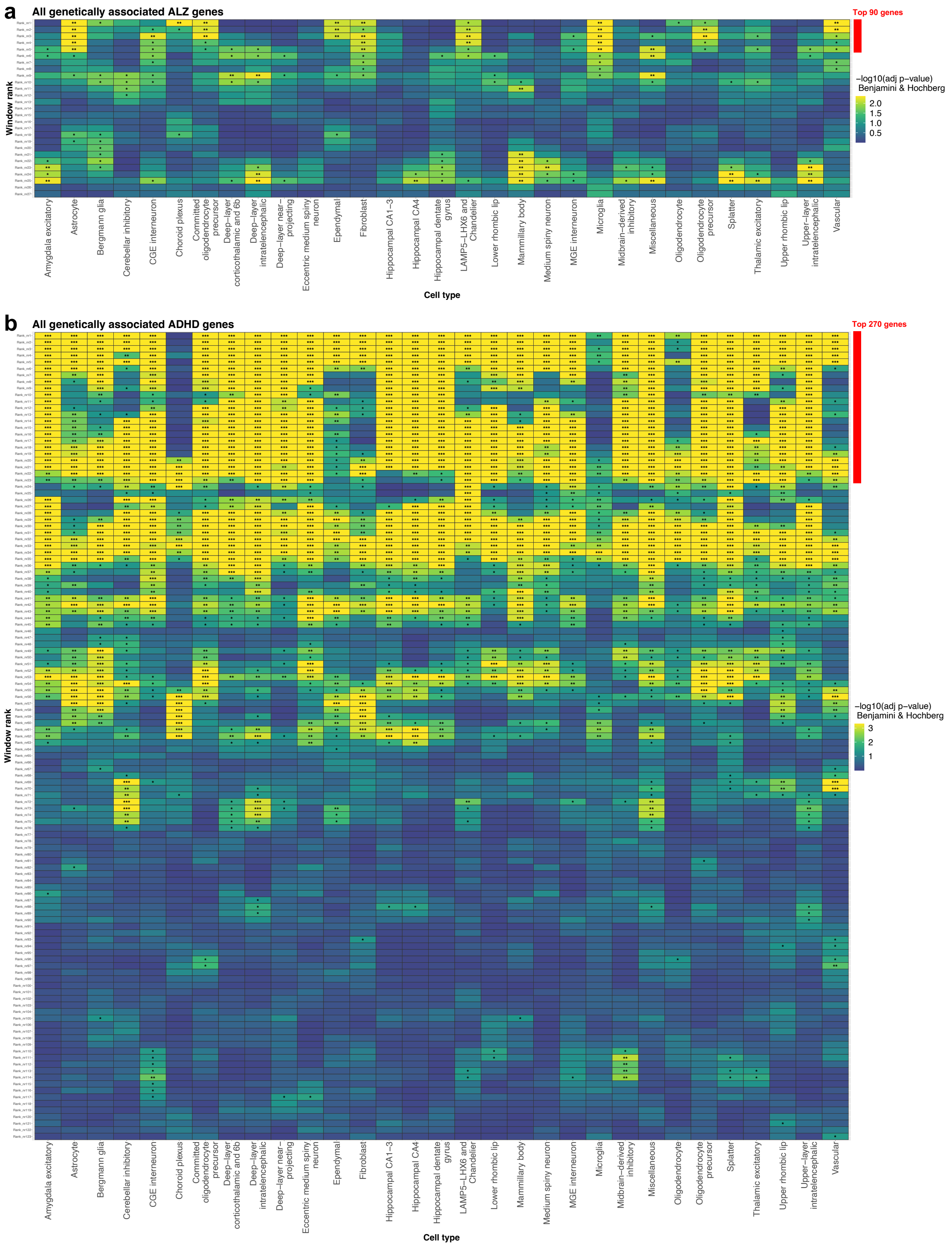


**Supplementary Figure 21. Enrichment of ALZ and ADHD across ranked gene lists in the adult human brain.** Trait enrichment with STEAM in cell types (using meta cells) for genetically associated ranked gene lists across superclusters in the adult human brain for **a)** ALZ, and **b)** ADHD. Each ranked widow consists of 50 genes, moving by 10 genes. Multiple testing correction with Benjamini & Hochberg, * $p < 0.05$, ** $p < 0.01$ and *** $p < 0.001$. Top genes selected with STEAM marked in red. CA = cornu Ammonis. MGE = Medial Ganglionic Eminence. CGE = Caudal Ganglionic Eminence.

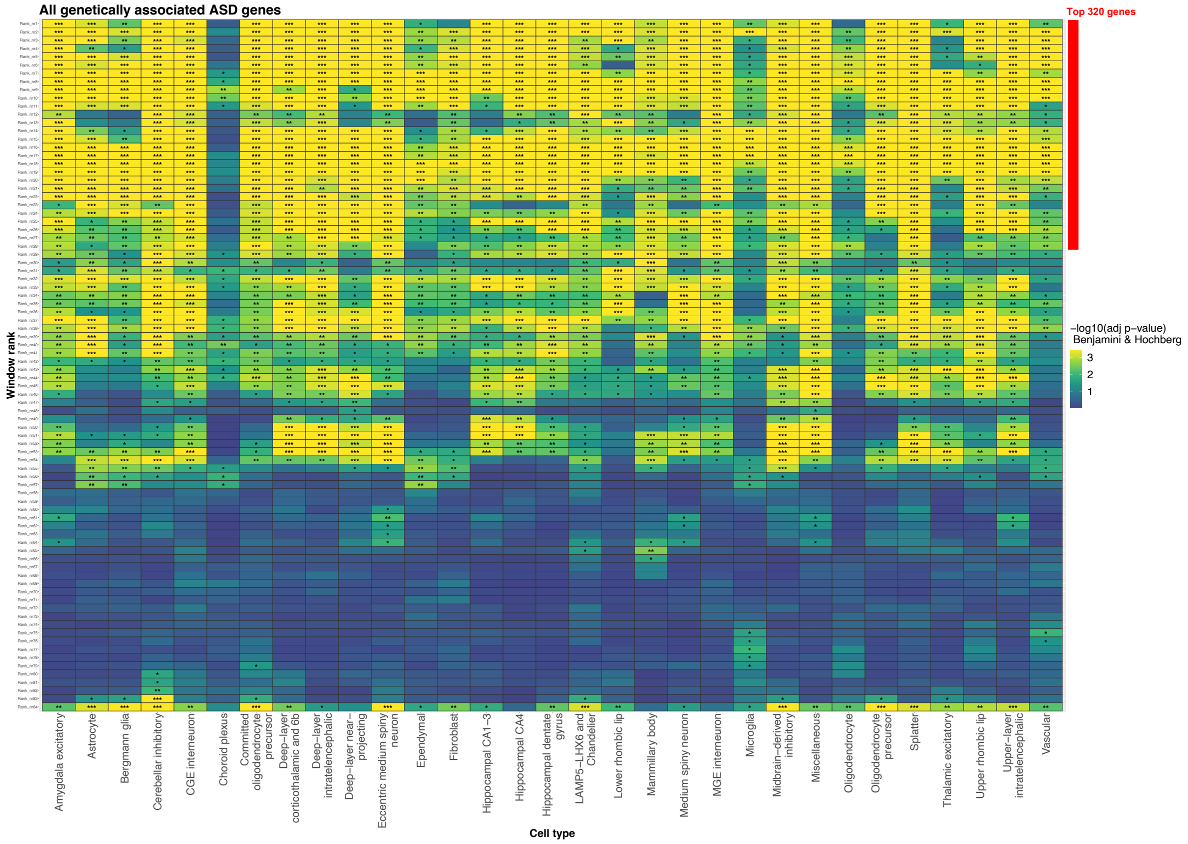


**Supplementary Figure 22. Enrichment of ASD across ranked gene lists in the adult human brain.** Trait enrichment with STEAM in cell types (using meta cells) for genetically associated ranked gene lists across superclusters in the adult human brain for ASD. Each ranked widow consists of 50 genes, moving by 10 genes. Multiple testing correction with Benjamini & Hochberg, * p < 0.05, ** p < 0.01 and *** p < 0.001. Top genes selected with STEAM marked in red. CA = cornu Ammonis. MGE = Medial Ganglionic Eminence. CGE = Caudal Ganglionic Eminence.

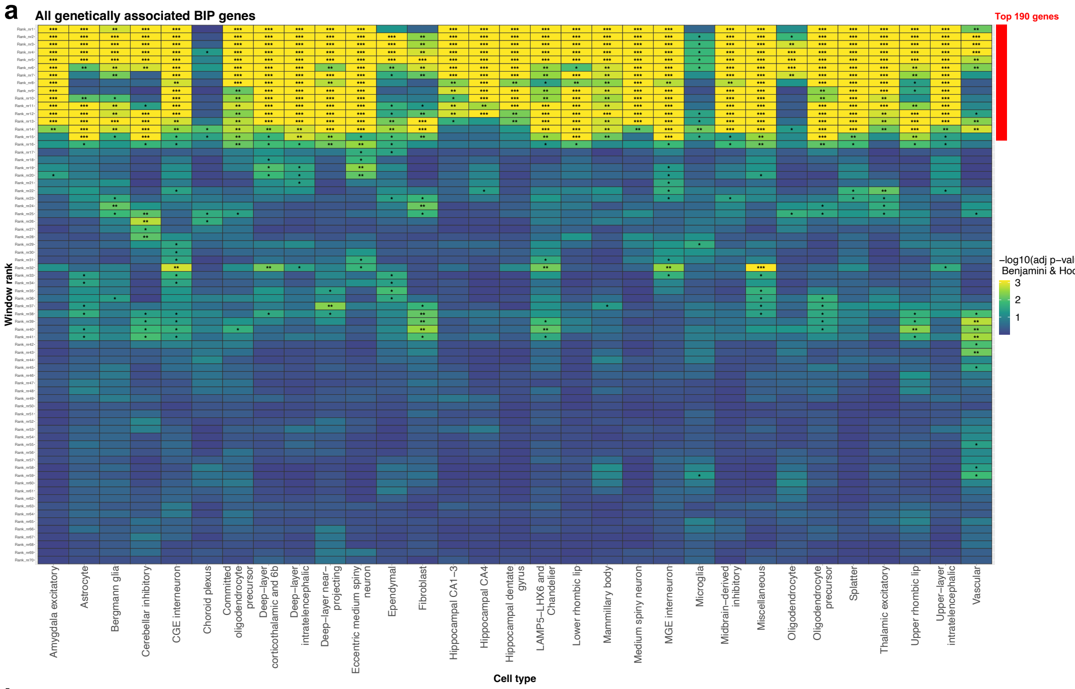
a
All genetically associated BIP genes
Top 190 genes
Window rank
−log10(adj p−value)
Benjamini & Hochberg
3
2
1
Amygdala excitatory
Astrocyte
Bergmann glia
Cerebellar inhibitory
CGE interneuron
Choroid plexus
Committed oligodendrocyte precursor
Deep−layer corticothalamic and 6b
Deep−layer intratelencephalic
Deep−layer near−projecting
Eccentric medium spiny neuron
Ependymal
Fibroblast
Hippocampal CA1−3
Hippocampal CA4
Hippocampal dentate gyrus
LAMP5−LHX6 and Chandelier
Lower rhombic lip
Mammillary body
Medium spiny neuron
MGE interneuron
Microglia
Midbrain−derived inhibitory
Miscellaneous
Oligodendrocyte
Oligodendrocyte precursor
Splatter
Thalamic excitatory
Upper rhombic lip
Upper−layer intratelencephalic
Vascular
Cell type

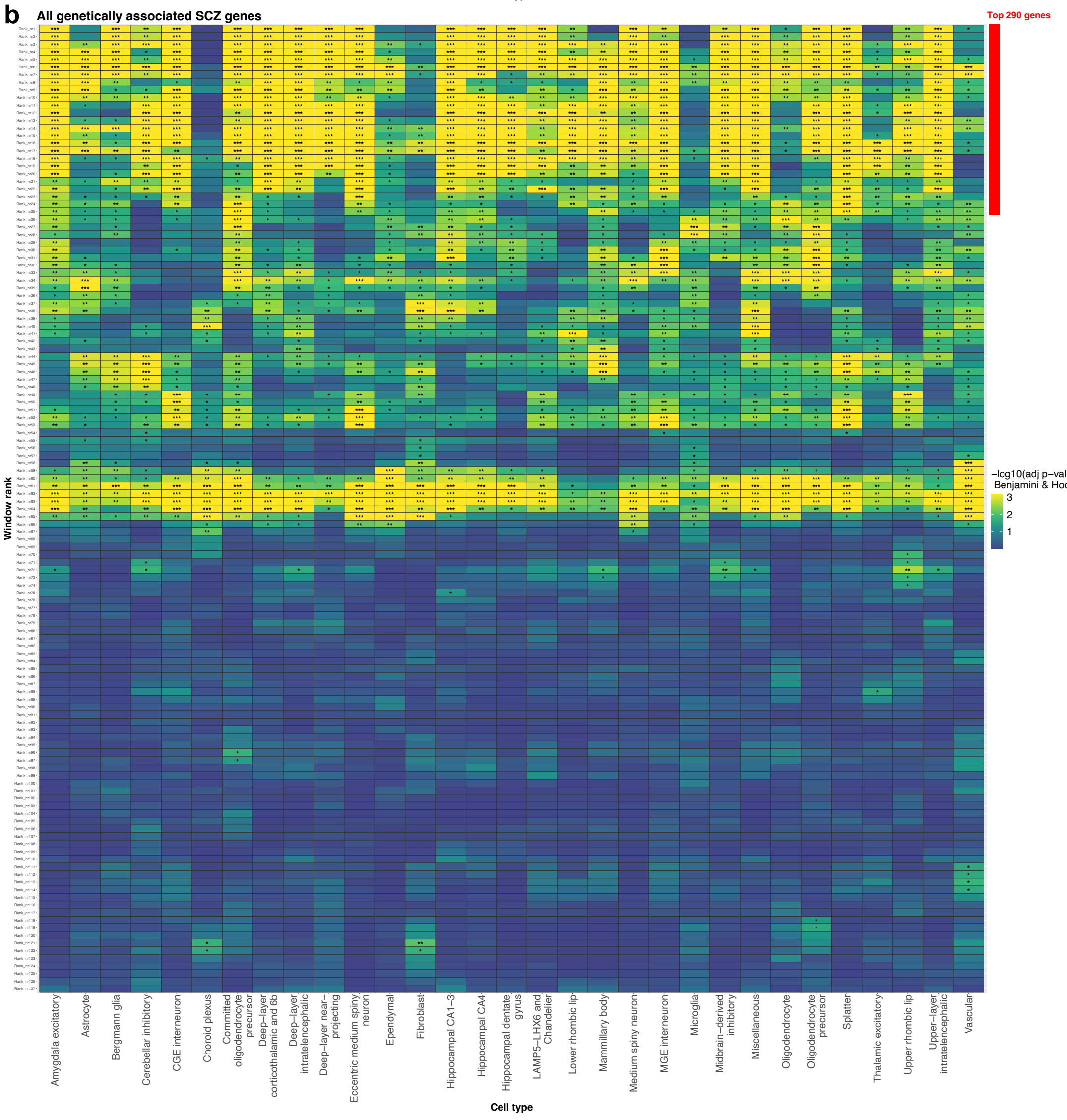
b
All genetically associated SCZ genes
Top 290 genes
Window rank
−log10(adj p−value)
Benjamini & Hochberg
3
2
1
Amygdala excitatory
Astrocyte
Bergmann glia
Cerebellar inhibitory
CGE interneuron
Choroid plexus
Committed oligodendrocyte precursor
Deep−layer corticothalamic and 6b
Deep−layer intratelencephalic
Deep−layer near−projecting
Eccentric medium spiny neuron
Ependymal
Fibroblast
Hippocampal CA1−3
Hippocampal CA4
Hippocampal dentate gyrus
LAMP5−LHX6 and Chandelier
Lower rhombic lip
Mammillary body
Medium spiny neuron
MGE interneuron
Microglia
Midbrain−derived inhibitory
Miscellaneous
Oligodendrocyte
Oligodendrocyte precursor
Splatter
Thalamic excitatory
Upper rhombic lip
Upper−layer intratelencephalic
Vascular
Cell type

**Supplementary Figure 23. Enrichment of BIP and SCZ across ranked gene lists in the adult human brain.** Trait enrichment with STEAM in cell types (using meta cells) for genetically associated ranked gene lists across superclusters in the adult human brain for **a)** BIP and **c)** SCZ. Each ranked widow consists of 50 genes, moving by 10 genes. Multiple testing correction with Benjamini & Hochberg, * $p < 0.05$, ** $p < 0.01$ and *** $p < 0.001$. Top genes selected with STEAM marked in red. CA = cornu Ammonis. MGE = Medial Ganglionic Eminence. CGE = Caudal Ganglionic Eminence.

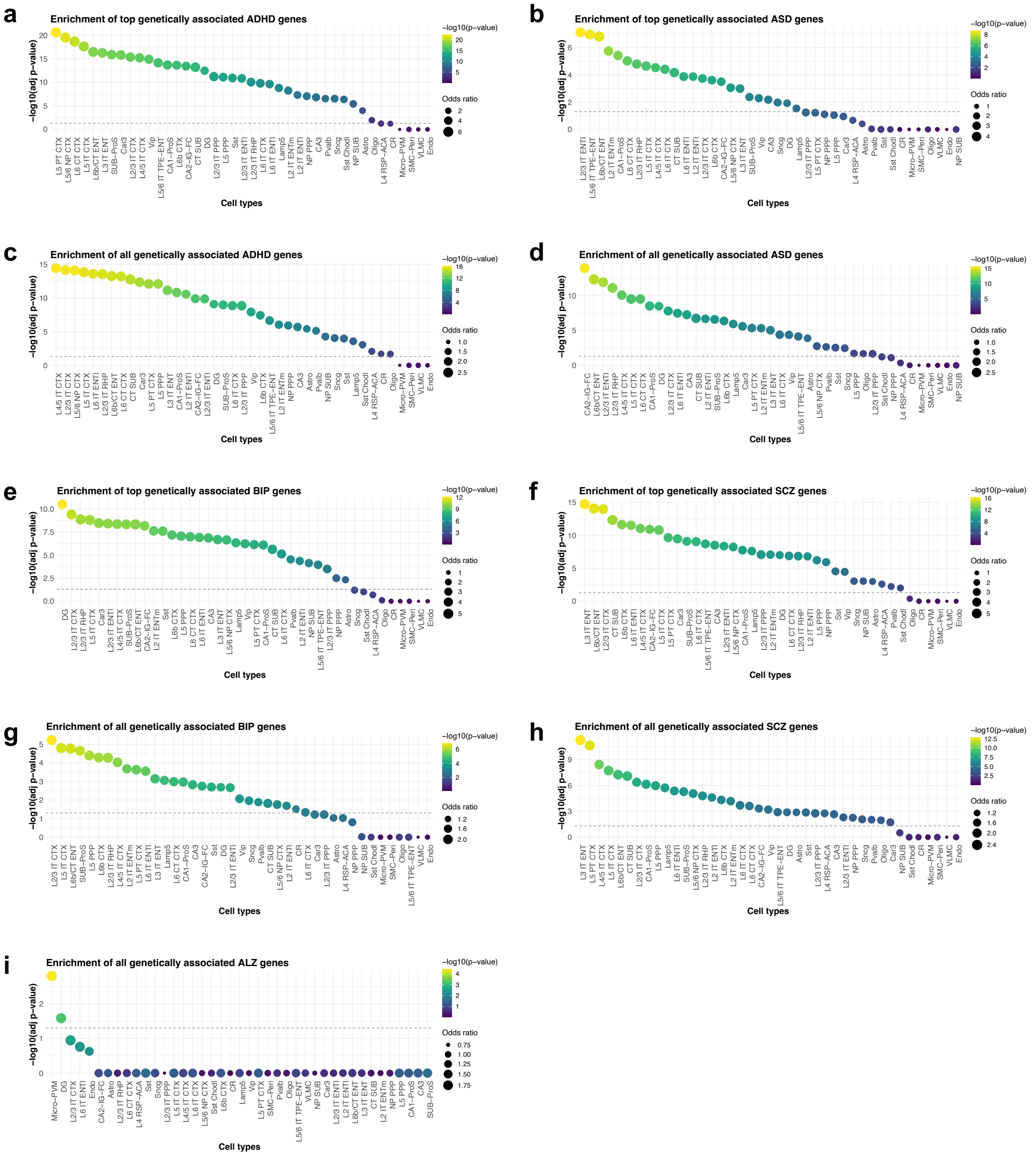


**Supplementary Figure 24. Trait enrichment in the mouse cortex and hippocampus.** Trait enrichment across cell types (using meta cells) for genetically associated genes (top genes selected with STEAM) among differentially expressed genes across subclasses in the mouse cortex and hippocampus. **a)** Top ADHD genes, **b)** top ASD genes, **c)** all ADHD genes, **d)** all ASD genes, **e)** top BIP genes, **f)** top SCZ genes, **g)** all BIP genes, **h)** all SCZ genes and **i)** all ALZ genes (grey dotted line, adj. p-value = 0.05). List of mouse cell subclass abbreviations can be found in Supplementary Table 6.

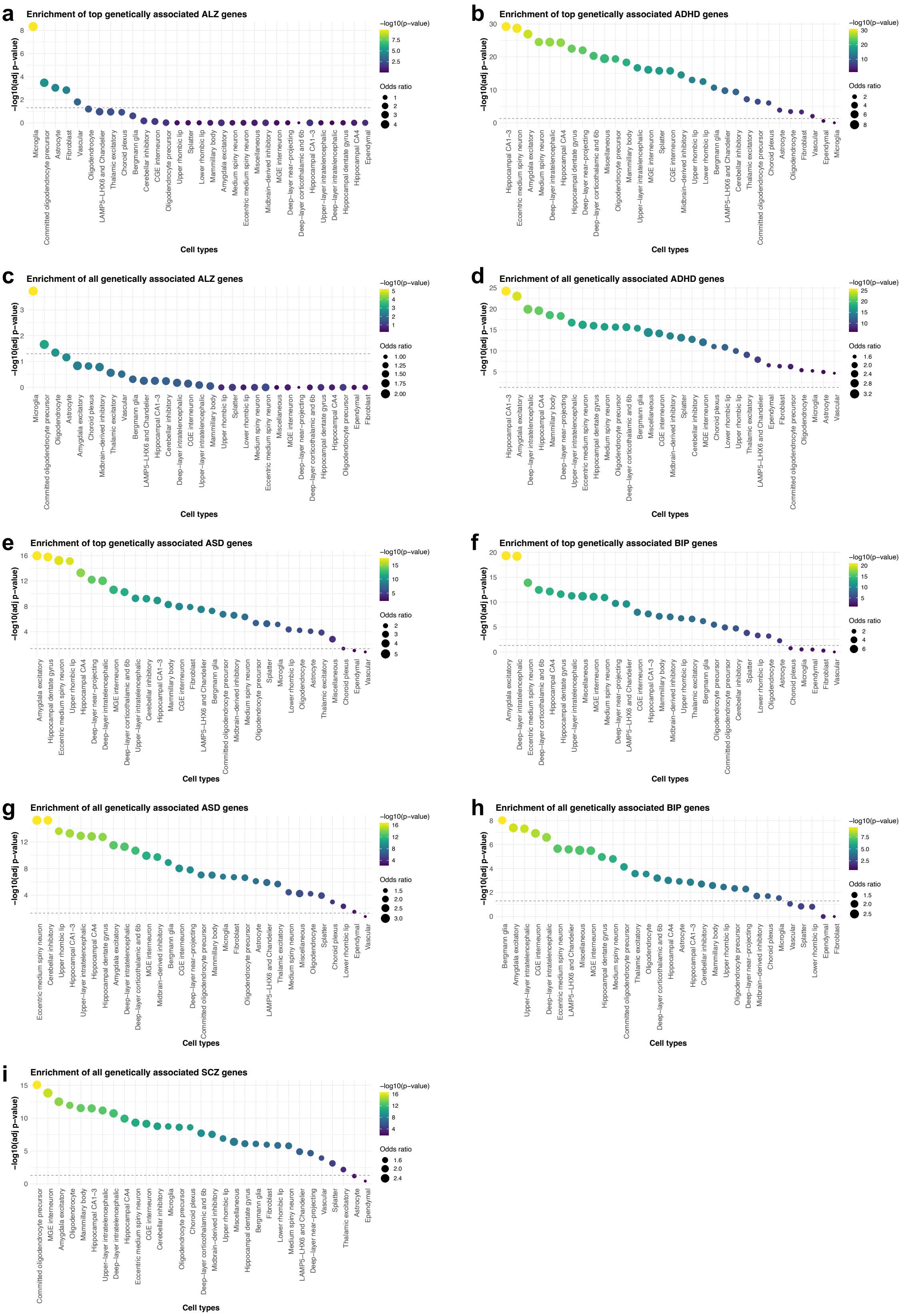


**Supplementary Figure 25. Trait enrichment in the adult human brain.** Trait enrichment across cell types (using meta cells) for genetically associated genes (top genes selected with STEAM) among differentially expressed genes across superclusters in the adult human brain. **a)** Top ALZ genes, **b)** top ADHD genes, **c)** all ALZ genes, **d)** all ADHD genes, **e)** top ASD genes, **f)** top BIP genes,

**g)** all ASD genes, **h)** all BIP genes and **i)** all SCZ genes (grey dotted line, adj. p-value = 0.05). CA = cornu Ammonis. MGE = Medial Ganglionic Eminence. CGE = Caudal Ganglionic Eminence.

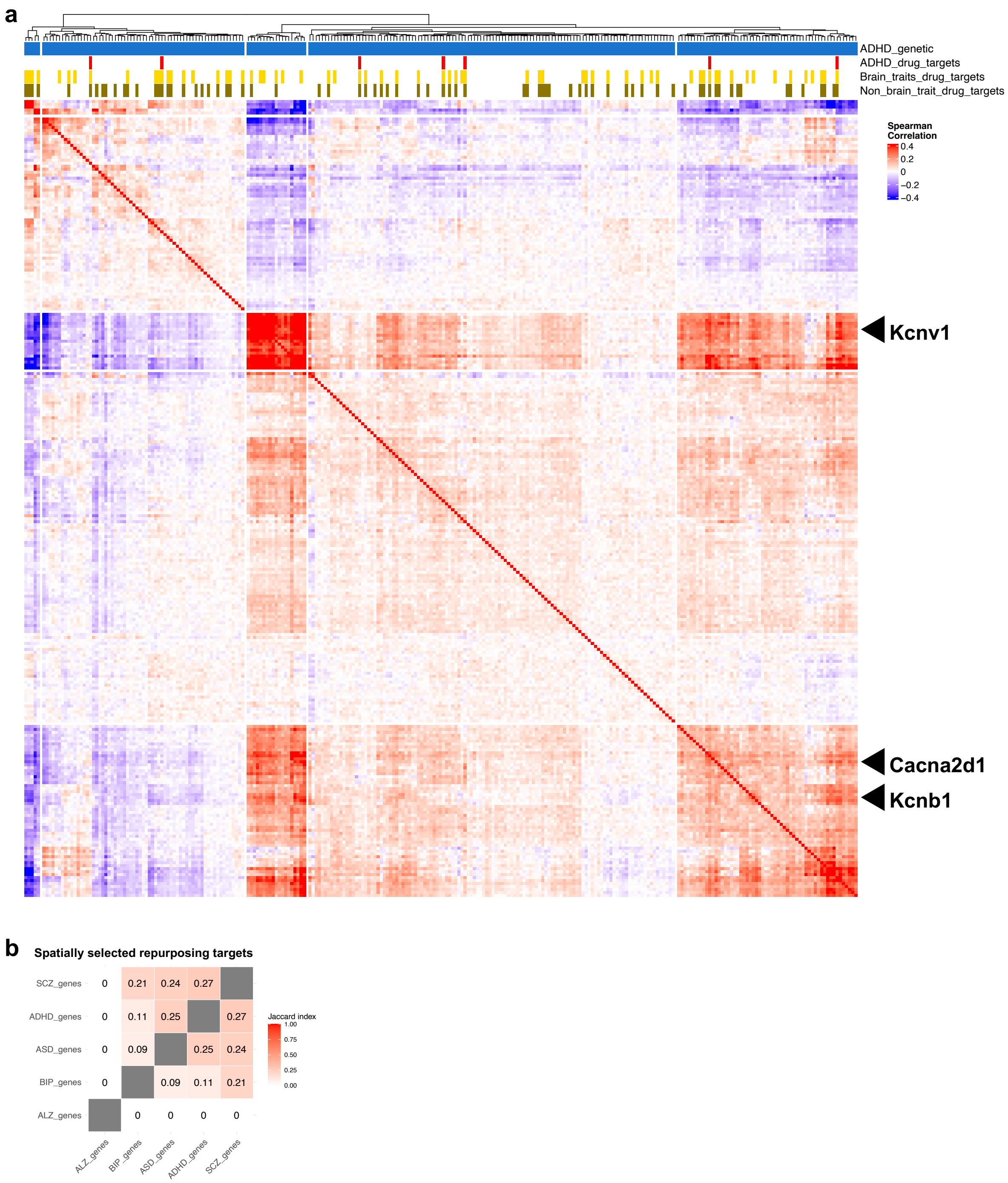


**Supplementary Figure 26. Spatial patterns of drug targets for ADHD. a)** Gene-gene spatial correlation heatmaps for the index mouse brain coronal tissue section, for genes genetically implicated in ADHD, with drug target status indicated. Arrows indicate drug targets of high interest shown in Figure 6 b. **b)** Overlap of trait-specific genetically associated drug target repurposing candidates.

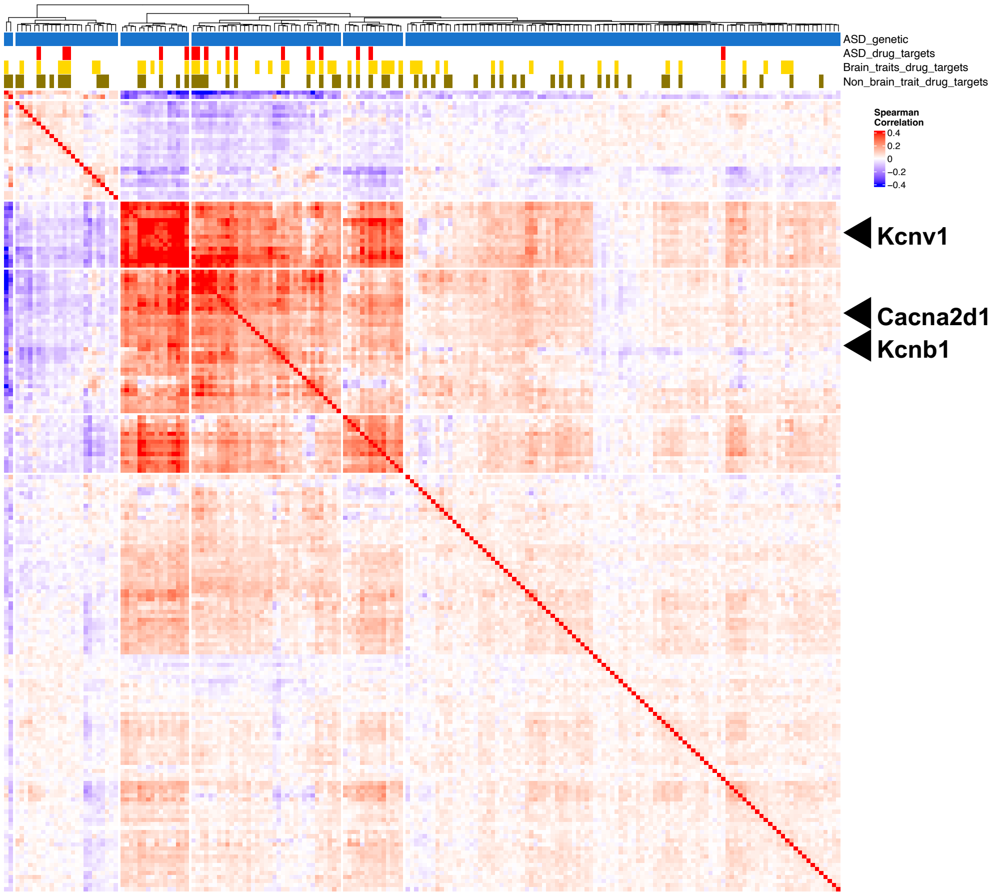


**Supplementary Figure 27. Spatial patterns of drug targets for ASD.** Gene-gene spatial correlation heatmaps for the index mouse brain coronal tissue section, for genes genetically implicated in ASD, with drug target status indicated. Arrows indicate drug targets of high interest shown in Figure 6 b.

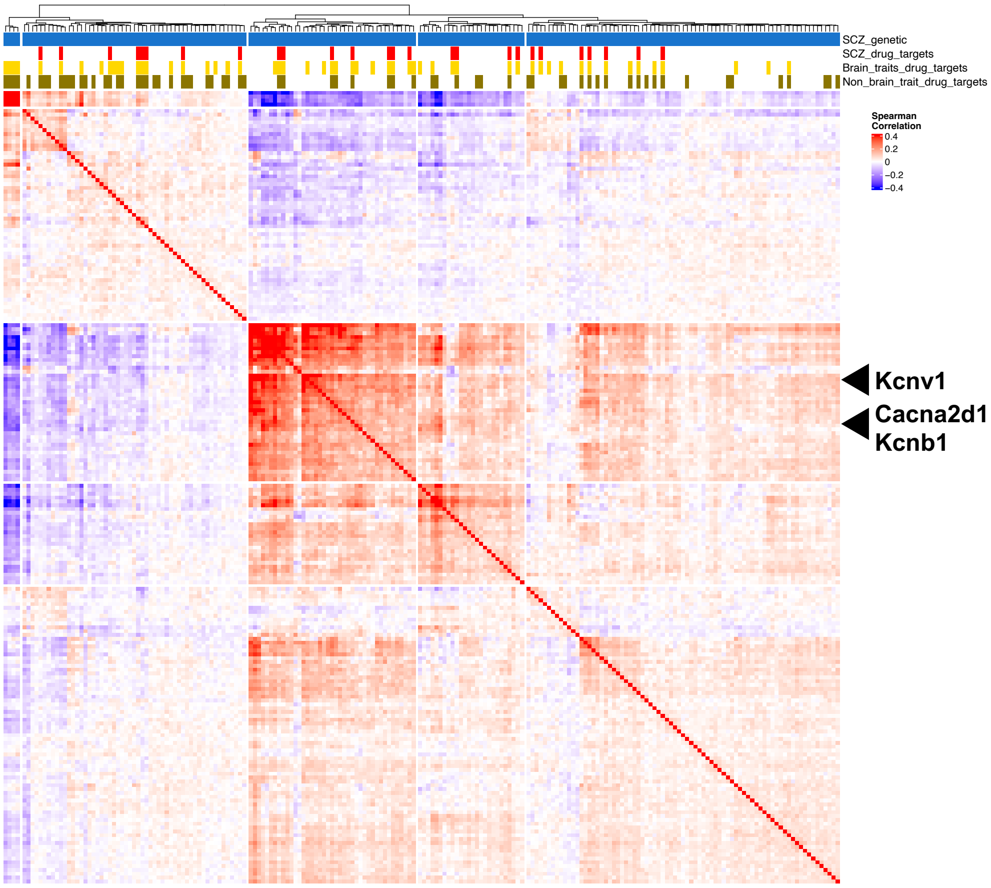


**Supplementary Figure 28. Spatial patterns of drug targets for SCZ.** Gene-gene spatial correlation heatmaps for the index mouse brain coronal tissue section, for genes genetically implicated in SCZ, with drug target status indicated. Arrows indicate drug targets of high interest shown in Figure 6 b.

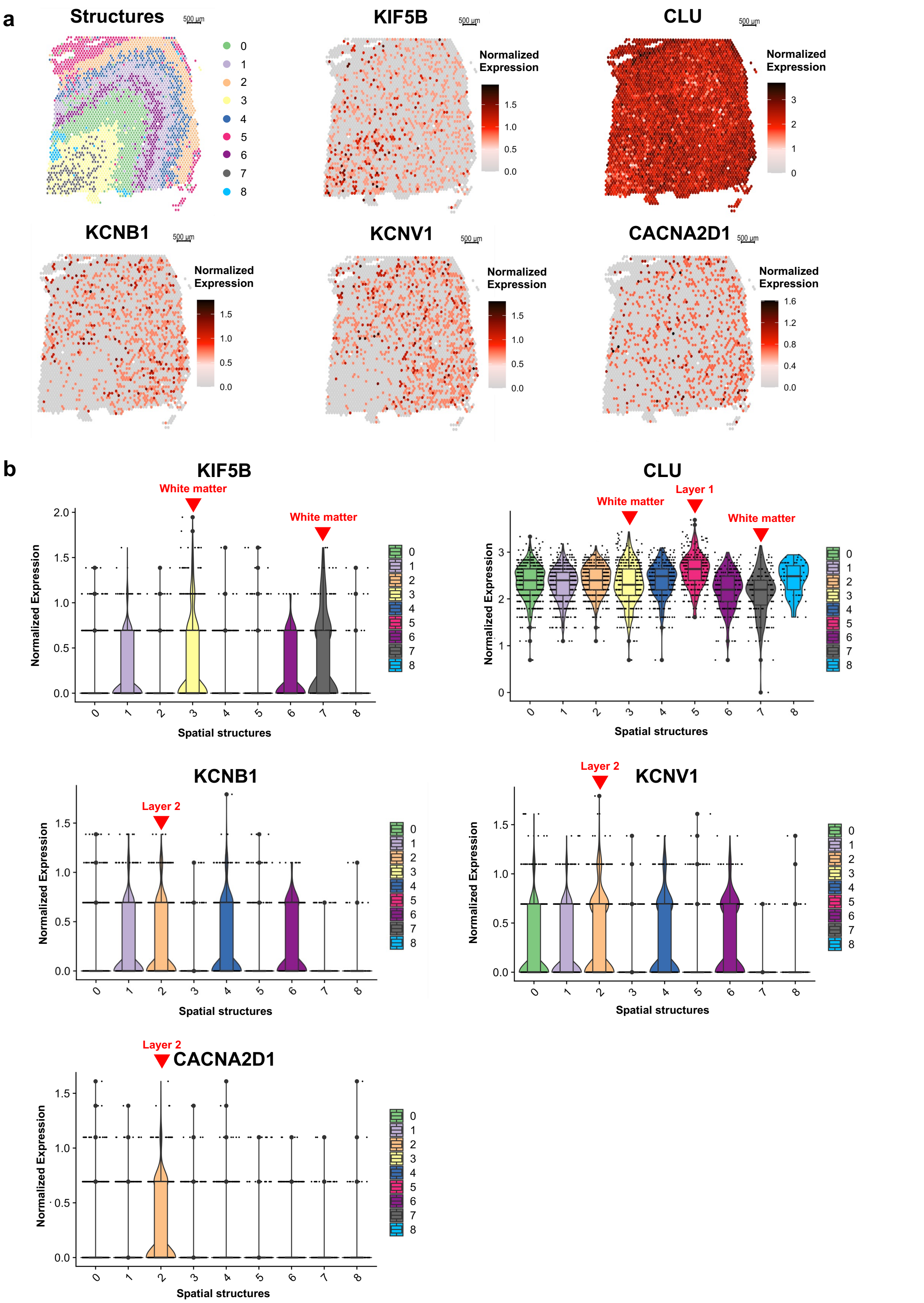


**Supplementary Figure 29. Putative drug target candidates in the human brain.** Normalized gene expression in the human cortex for dataset (brain_human_151673) for a select set of putative drug target candidates. **a)** Spatial visualization of normalized expression. **b)** Structure specific normalized expression. Arrows and text (red) indicating relevant tissue structures. Scale bar 500µm.

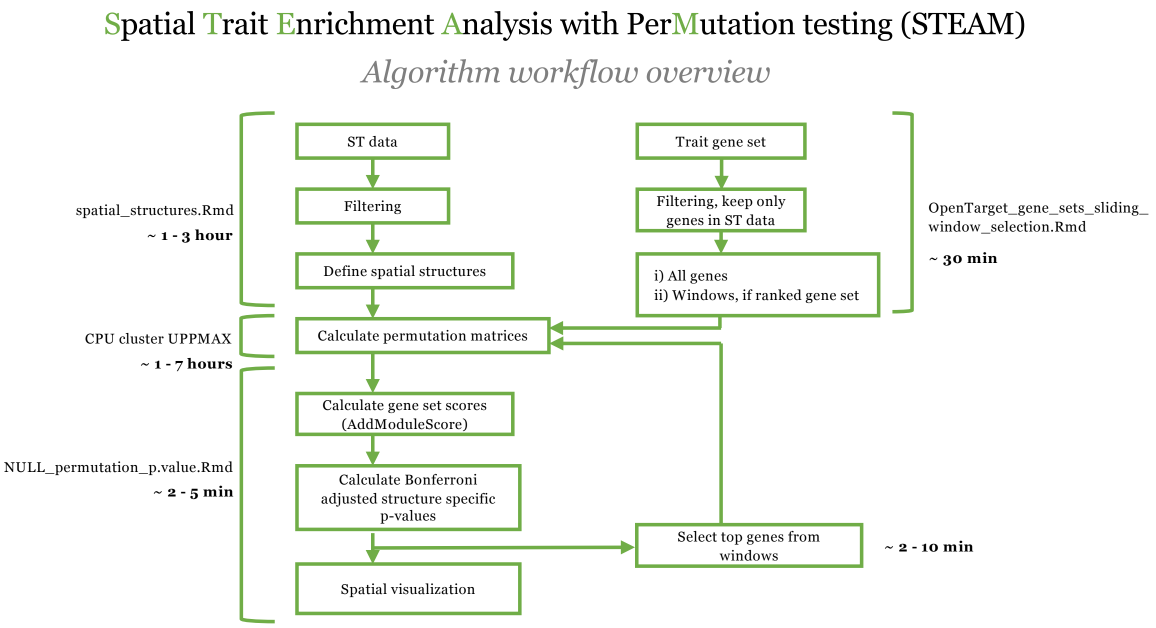


**Supplementary Figure 30. STEAM workflow.** Computational algorithm workflow overview, including estimated time for analysis.

### Supplementary Table texts

**Supplementary Table 1. Spatial transcriptomics datasets.** Links to published spatial transcriptomics datasets, tissue, species, library preparation method and dataset name within this publication.

**Supplementary Table 2. Spatial structure annotation.** Spatial barcodes and seurat cluster number for all datasets.

**Supplementary Table 3. Differentially expressed genes.** Across spatial seurat clusters for all datasets.

**Supplementary Table 4. Open Targets gene list.** Trait names, trait ID, number of genes with genetic and drug associations, Open Targets release version of each gene list and Open Targets trait ID.

**Supplementary Table 5. STEAM overview and results.** Dataset and trait-specific information and results. For both selected and non-selected trait-tissue pairs. Bonferroni correction.

**Supplementary Table 6. STEAM results across ranked gene lists for sc/snRNA-seq.** Trait enrichments for ALZ, ADHD, ASD, BIP and SCZ, using the Allen Brain Atlas of the mouse cortex and hippocampus and a human brain dataset. Full list of mouse cell subclass abbreviations. Benjamini & Hochberg correction.

**Supplementary Table 7. Complementary enrichment results for sc/snRNA-seq.** Trait enrichments for all and top genetically associated genes for ALZ, ADHD, ASD, BIP and SCZ, using the Allen Brain Atlas of the mouse cortex and hippocampus and a human brain dataset. Bonferroni correction.

**Supplementary Table 8. Drug repurposing candidates.** Candidate genes and drug gene list overlap with other study traits, for ALZ, ADHD, ASD, BIP and SCZ, based on spatially informed correlation analysis in the mouse coronal dataset.
